# Spatiotemporal Profiling Unveiling the Cellular Organization Patterns and Local Protumoral Immune Microenvironment Remodeling in Early Lung Adenocarcinoma Progression

**DOI:** 10.1101/2023.10.29.564580

**Authors:** Haojie Chen, Yizhou Peng, Yana Li, Qiang Zheng, Yue Zhao, Pengcheng Liu, Zhigang Wu, Yang Wo, Hui Hong, Yihua Sun, Zhen Shao

## Abstract

Spatial cellular organization patterns (COPs) in tumor microenvironment influence the tumor progression and therapeutic response, however, little is known about the cellular composition and functional potential of these multicellular structures during lung adenocarcinoma progression. Here, we integrate spatial transcriptomics with single cell RNA sequencing to characterize the local tumor and immunological landscape of samples from 8 patients with early-stage lung adenocarcinoma at different pathological stages. We identified ten COPs that show distinct associations with local immune states and clinical outcomes, including survival and therapy response. The local infiltration levels of regulatory and dysfunctional immune cells are increased with pathological progression. Cell-to-cell interactions between malignant cells and tumor microenvironment (TME) cells were involved in protumor immune state remodeling. Finally, we detected a group of malignant cells that were specifically located at the tumor boundary, representing a more aggressive state, were involved in the invasion of invasive adenocarcinoma (IAC). Altogether, these results can improve our understanding of the local microenvironment characteristics that underlie LUAD progression and may facilitate the identification of drug targets to prevent invasive progression and biomarkers for diagnosis.

## Introduction

Lung adenocarcinoma (LUAD) is the most prevalent subtype of lung cancer and accounts for most cancer-related fatalities in both males and females^1^. LUAD can be classified into two main types: minimally invasive adenocarcinoma (MIA) and invasive adenocarcinoma (IAC). Compared to MIA, IAC, in particular, is characterized by aggressive histopathological features, such as invasive growth patterns where cancer cells invade the surrounding lung tissue^2^. Previous studies have depicted the epigenetic, transcriptomic and metabolomic dynamics with LUAD pathological progression using bulk sequencing technologies and revealed critical molecular alterations underlying the initiation and progression of LUAD^3–6^. However, our understanding of the remodeling of the tumor microenvironment (TME) and the specific characteristics of different cell types within this context remains limited.

In recent years, single cell RNA sequencing (scRNA-seq) has emerged as a valuable tool for investigating the molecular and cellular heterogeneity within the TME of LUAD. These studies demonstrated that the enrichment of regulatory T cells and decrease in cytotoxicity in CD8 T cells were related to progression from MIA to IAC^7,8^. Notably, tissue dissociation leads to loss of spatial information, which plays a crucial role in determining cellular functionality and higher order cellular organization in the TME. To overcome this challenge, spatial transcriptomics (STs) was developed as a technique that simultaneously captures the location and sequence of each transcript within an intact tissue^9^. However, integrating multiple STs data from different LUAD patients to discover recurrent cellular organization patterns and characterize functional states of these spatially organized cell types remains a significant hurdle^10^.

To address these issues, scRNA-seq data and STs data from patients in different progression stages were generated and integrated to create a comprehensive spatial map of early-stage LUAD to investigate the landscape of spatial cellular organization patterns (COPs) during LUAD progression and its relationship to tumor immune microenvironment (TIME) remodeling. Thus, further reveal mechanistic insights into disease progression and unveil biomarkers of clinical prognosis and therapeutic responses. We identified ten COPs, including COP enriched with normal epithelial cells, COPs enriched with malignant cells, COP colocalized with lymphoid aggregates (LAs) and tertiary lymphoid node structures (TLSs) and COP as a macrophage niche. These COPs exhibited distinct associations with immune states and clinical outcomes. In addition, we discovered a group of malignant cells with invasive and immune suppressive potential that were specifically located at the tumor boundary. Overall, our study unravels local protumor remodeling and immunosuppression that occur during the pathological progression of LUAD, as well as spatially organized cellular structures in the LUAD ecosystem that can predict patient clinical outcomes. The findings of this study will provide valuable theoretical evidences for improving clinical prognosis and furthering our understanding of the relationship between spatial cellular organization and its pathological progression.

## Results

### Obtaining STs data and scRNA-seq data from early-stage LUAD patients at different pathological stages

In order to generate a comprehensive spatial map of malignant and immune cells in early-stage LUAD during histopathological progression, we obtained tissue specimens from eight early-stage LUAD patients including four cases of MIA and four cases of IAC and performed ST sequencing using the 10X Genomics Visium Platform. However, this spatial method is unable to create a single-cell resolution spatial transcriptomic map, which underscores the need to integrate STs data with scRNA-seq data to maximize the resolution. Thus, we further performed scRNA-seq on 11 tissue specimens from the corresponding patients, composed of eight samples from tumor tissues and three samples from matched normal lung tissues. The bulk normal lung and tumor tissues of all patients were also used for RNA-seq and whole-exome sequencing (WES) (Figure 1A and Supplementary Table 1).

**Figure 1.**
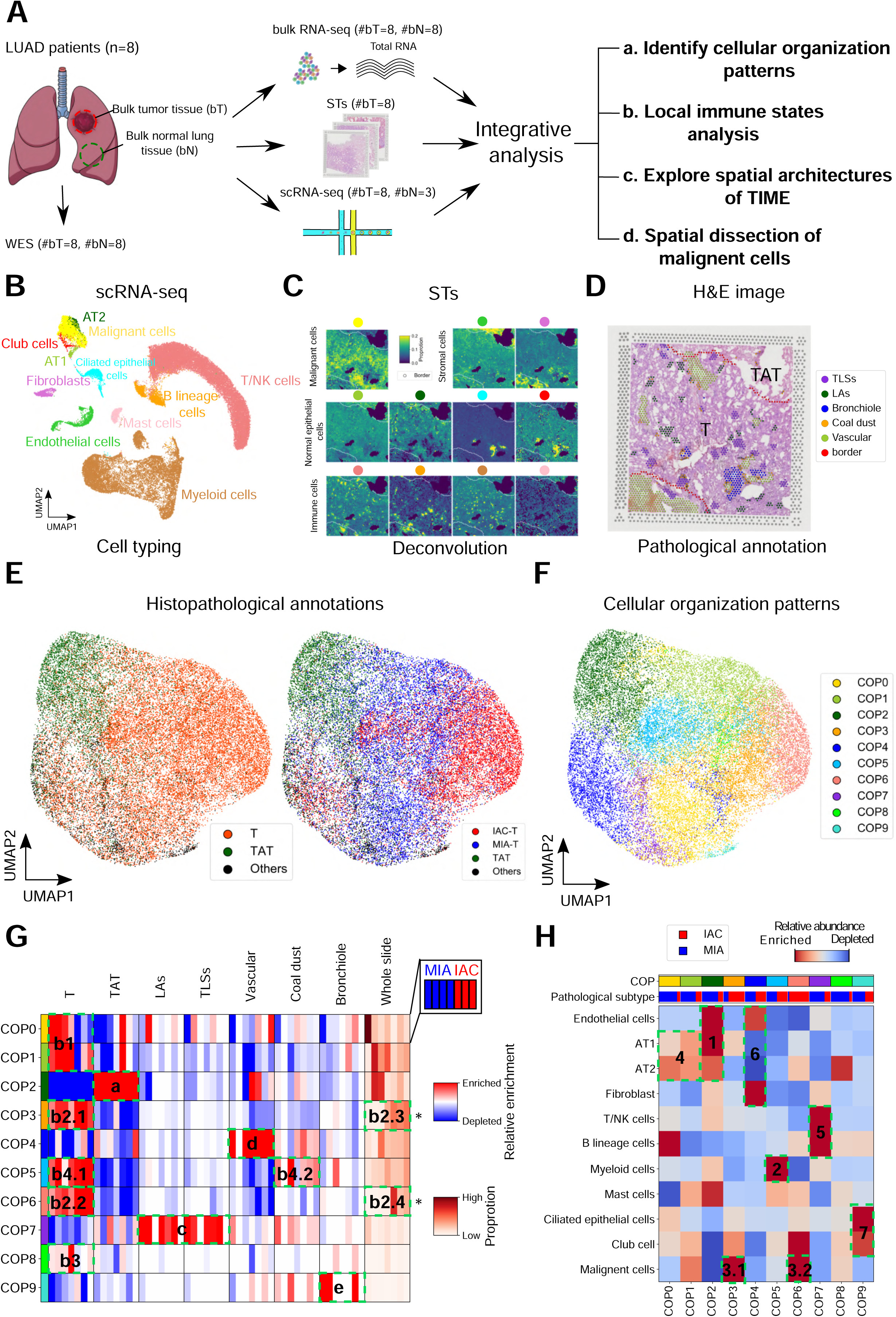
Defining the landscape of cellular organization patterns in LUAD pathological progression by integrating Spatial Transcriptomics with single cell RNA-seq. **(A)** Schematic depicting acquisition of whole exome sequencing (WES), bulk RNA-seq, spatial transcriptomics (STs) and single-cell RNA-seq (scRNA-seq) data from eight LUAD patients with MIA and IAC, integration of STs data with scRNA-seq data identifies cellular organization patterns (COPs) in LUAD, reveals the local immune states during LUAD progression, explores the role of spatial architectures of immune cells and dissects spatial characteristics of malignant cells. **(B)** UMAP view of 11 cell types identified from scRNA-seq data. **(C)** Location and proportion of 11 cell types in a representative slide (P01). **(D)** HE image showing the histological annotation of a representative slide (P01). **(E)** UMAP showing STs spots colored by histopathological annotation types. T, tumor tissues, TAT, tumor adjacent tissues. **(F)** UMAP of ten COPs discovered in LUAD by integrating STs data with scRNA-seq data. **(G)** Heatmap depicting enrichment/depletion level of each COP in each histological/pathological region and the proportion in each patient. ‘*’ indicates rank sum test derived *p*-value less than 0.05. **(H)** Heatmap of relative cellular compositions in each COP.

In STs data, after basic quality control, we obtained 31,666 spots for further analysis with a median of 13,089 unique molecular identifiers (UMIs) and 4,332 features per spot (Supplementary Figure 1; Details see Methods). To assess the concordance between the transcriptomic characteristics and the hand-labeled pathological annotations, we first employed a signature score-based strategy to evaluate the enrichment degree of tumor upregulated genes and normal lung upregulated genes in each spot. The tumor upregulated genes and normal lung upregulated genes were identified through differentially expressed genes (DEGs) analysis in bulk RNA-seq data (Supplementary Figure 2). Next, we evaluated the performance of these two signature scores in distinguishing cancerous and noncancerous spots on each slide. The classification based on two signature scores yielded high area under the curve (AUC) values, with a median AUC of 0.84. In contrast, classification using randomly selected genes resulted in much lower AUC values, with a median AUC of 0.42. This consistent performance was observed across most of slides, except P12 (Supplementary Figure 3). According to the pathologists’ assessment, the distinction between tumor tissue and adjacent tumor tissue was not clear in the HE image of this particular patient. Hence, slide P12 was excluded from downstream analyses.

### Identifying distinct cell types in the LUAD TME

To investigate the spatial architecture of LUAD and address the resolution limitation of this spatial method, we harvested 67,083 cells using 10X scRNA-seq^11^ from 11 biopsy specimens of normal lung tissues (n=3), MIA tumor tissues (n=4) and IAC tumor tissues (n=4). After filtering, a total number of 44,003 cells were retained for subsequent analyses with a median of 4,791 unique molecular identifiers (UMIs) count and 1,665 features per cell (Supplementary Figure 4; Details see Methods). Finally, these cells were clustered into eight major cellular lineages or cell types, which were annotated based on key marker gene expression, comprising epithelial types (epithelial cells and ciliated epithelial cells), stromal types (endothelial cells and fibroblasts) and immune types (T/NK cells, B lineage cells, myeloid cells and mast cells) (Supplementary Figure 5A-B and Supplementary Table 2). Importantly, each cell type was found to be present in multiple samples and tissue types, indicating no strong batch effect in the scRNA-seq data. Notably, T/NK cells and myeloid cells represented a significant proportion of the cell population, suggesting an opportunity to further elucidate the immune composition of the LUAD TIME (Supplementary Figure 5C).

Given previous studies suggesting that epithelial cells are the cell of origin of LUAD, we took the analysis further by dividing epithelial cells from all samples into 10 clusters to identify malignant cells (Supplementary Figure 5D). By examining the expression of established normal epithelial markers, we successfully identified populations corresponding to alveolar type I (AT1, sc-epi-C7, *AGER*), alveolar type II (AT2, sc-epi-C8, *SFTPC*), and club cell (sc-epi-C9, *PIGR*). Meanwhile, other epithelial sub-populations (i.e., sc-epi-C[0-6]) may represent tumor epithelial cells, as most of them highly expressed tumor markers, such as *CEACAM5* (Supplementary Figure 5E and Supplementary Table 3). To differentiate malignant cells from non-malignant ones, *inferCNV*^12^ was utilized to estimate copy number variations (CNVs) from scRNA-seq data in each epithelial cell. Subsequently, a CNV score was calculated to quantify the CNV level (Details see Methods). The cells in sc-epi-C[0-6] exhibited overall increased CNV scores, which further supported that these clusters predominantly consisted of malignant cells (Supplementary Figure 5F). We also observed a gradually increase in CNV scores across the stages of pathological progression (Supplementary Figure 5G). Altogether, these results represented transcriptomic and genetic evidences supporting sc-epi-C[0-6] as malignant cells in LUAD. Finally, utilizing the scRNA-seq data, we defined 11 major cell types in scRNA-seq data, which were then used to estimate the proportion of each cell type for a given capture spot (Figure 1B).

### Distinct cellular organization patterns of LUAD pathological progression related spatial heterogeneity

The spatial organization of various cell types within LUAD is believed to play a crucial role in cancer progression^13^. Notably, high-order arrangements of cells, such as lymphocyte aggregates (LAs) and tertiary lymphoid structures (TLSs), may represent structural and functional components in tumor tissues^14,15^. To identify and analyze these spatial patterns, we employed *SPOTlight*^16^ to deconvolute the cellular composition of each spot in each slide (Figure 1C), and created a two-dimensional embedding of spotwise cellular compositions using uniform manifold approximation and projection (UMAP). Spots with similar cell type compositions were clustered closely to each other in this UMAP space.

Next, we used unsupervised clustering to identify cellular organization patterns (COPs), which were then mapped onto pathologist annotated regions, allowing for relating well-known pathological and histological tissues to COPs defined by STs analysis (Figure 1D and Supplementary Figure 6; Details see Methods). We performed this analysis to integrate slides from different patients and the shared embedding space revealed a clear separation of cellular composition in both tumor (T) spots and tumor adjacent tissue (TAT) spots. Importantly, we observed distinct cellular compositions between spots from different pathological stages, indicating that a global cellular composition changes during pathological progression.

Using this approach, we totally identified ten COPs that delineated both known and new tissue architectures. Among these, COP2 was found to be specifically located in tumor adjacent tissues (TAT, Figure 1G, box a) and was also enriched for alveolar cells (AT1 and AT2) and endothelial cells (Figure 1H, box 1). In contrast, COP0, COP1, COP3, COP5, COP6 and COP8 were mainly enriched in tumor tissues (Figure 1G, box b[1-4]), but COP5 contains more spots annotated by coal dust than other COPs (Figure 1G, box b4.2) and was highly enriched for myeloid cells (Figure 1H, box 2), COP8 was a special case, which is annotated as a papillary histological subtype in only P13 (Figure 1G, box b3 and Supplementary Figure 7). Furthermore, we observed that COP0 and COP1 were highly enriched in tumor tissues of patients with MIA (Figure 1G, box b1), while the proportions of COP3 and COP6 are significantly increased in patients with IAC (Figure 1G, box b2.3 and box b2.4) and were highly enriched for malignant cells (Figure 1H, box 3.1 and box 3.2). By contrast, COP0 and COP1 are enriched for AT1 and AT2 (Figure 1H, box 4). COP7 was enriched in LAs or TLSs, Figure 1G, box c) and was enriched for T/NK cells and B cells (Figure 1H, box5), in line with recent report that B and T cell populations make up the bulk of TLSs-associated immune cells^14^. COP4 was enriched in vascular niche (Figure 1G, box d), and highly enriched for endothelial cells and fibroblasts (Figure 1H, box 6). COP9 is mostly located in bronchioles (Figure 1G, box e) and is enriched for bronchiole epithelial cells^17^, including ciliated epithelial cells and club cells (Figure 1H, box 7). Together, we summarized the COPs in LUAD and found that the proportions of tumor tissue enriched COPs (i.e., COP0, COP1, COP3 and COP6) are associated with pathological progression. We also identified COPs related to well-organized immune cell structures, such as LAs and TLSs. These results provide valuable insights into the spatial organization of different cell types in the LUAD tumor microenvironment.

### Building a comprehensive immune cell reference in the LUAD TIME

By utilizing multicellular resolution spatial transcriptomic profiling and previous defined COPs, we are able to quantify the immune cells surrounding malignant/normal epithelial cells at different progression stages. This allows us to evaluate the cellular states and functional potential of immune cells (i.e., functional or dysfunctional immune cells) that are either enriched or depleted in these areas, providing insight into the local immune state of these spots or COPs (Figure 2A). To this end, we proceeded to further divide the major immune cell types into 30 distinct clusters. These clusters include 6 CD4+ T cell clusters, 6 CD8+ T cell clusters, 5 natural killer cell (NK cell) clusters, 5 dendritic cell (DC) clusters, 2 macrophage clusters, 1 B cell cluster, 1 plasma cell (PC) cluster, 2 monocyte clusters, 1 neutrophil cluster, and 1 mast cell cluster (Figure 2B). The immune cell clusters were manually annotated by canonical markers and marker genes that highly expressed in each specific cluster (Figure 2B, Supplementary Figure 8 and Supplementary Table 4; Details see Methods). For instance, in the case of CD4+ T cells, regulatory T cells (Tregs) exhibit high expression of *FOXP3* and immune inhibitory receptors such as *CTLA4* and *TIGIT*. On the other hand, naive CD4+ T cells, central memory CD4+ T cells, and effector memory CD4+ T cells express *SELL*, *CD28*, and *BATF*, respectively. CD8+ T cells in an exhausted state were identified by the expression of *HAVCR2* (*TIM3*) and *LAYN* marker genes. Additionally, three clusters of CD8+ T cells (namely *GIMAP7*+ CD8+ effector T cells, *HSP1B1*+ CD8+ effector T cells, and *GZMK*+ CD8+ effector T cells) were found to exhibit high expression of cytotoxic-associated genes (*GZMA*/*B*) and *INFG*^18^, thus defining them as effector CD8+ T cells. Five distinct subpopulations of NK cells were identified and characterized based on their marker genes. One particular cluster of NK cells exhibited marker expression consistent with that of natural killer T cells (NKT)^19^, including NK cell markers such as *KIR2DL3* and *KLRF1*, as well as moderate expression of T cell markers including *CD3D*/*E*/*G*. As a result, this specific NK cell cluster was classified as an NKT cell subpopulation. Furthermore, five distinct dendritic cell subpopulations were identified and annotated by previously defined markers, including classical type 2 dendritic cells (*CLEC10A*), plasmacytoid dendritic cells (pDC, *LILRA4*), mature dendritic cells (*CCR7*), classical type 1 dendritic cells (*CLEC9A*) and Langerhans cells (*S100B*)^20^. Macrophages were separated into two subtypes, alveolar macrophages (AMs) and monocyte-derived macrophages (MDMs) with *MARCO* and *SPP1* as the respective marker genes^21^. Plasma cells exclusively expressed immunoglobulins such as *IGHA1* and *IGHG1*, enabling them to be distinguished from B cells expressing *MS4A1* (Figure 2B). In summary, we built a comprehensive reference of immune cell types present in the TIME of LUAD, which can be used for deconvolution to quantify the immune neighborhood of malignant/normal epithelial cells and reveal the local immune states.

**Figure 2.**
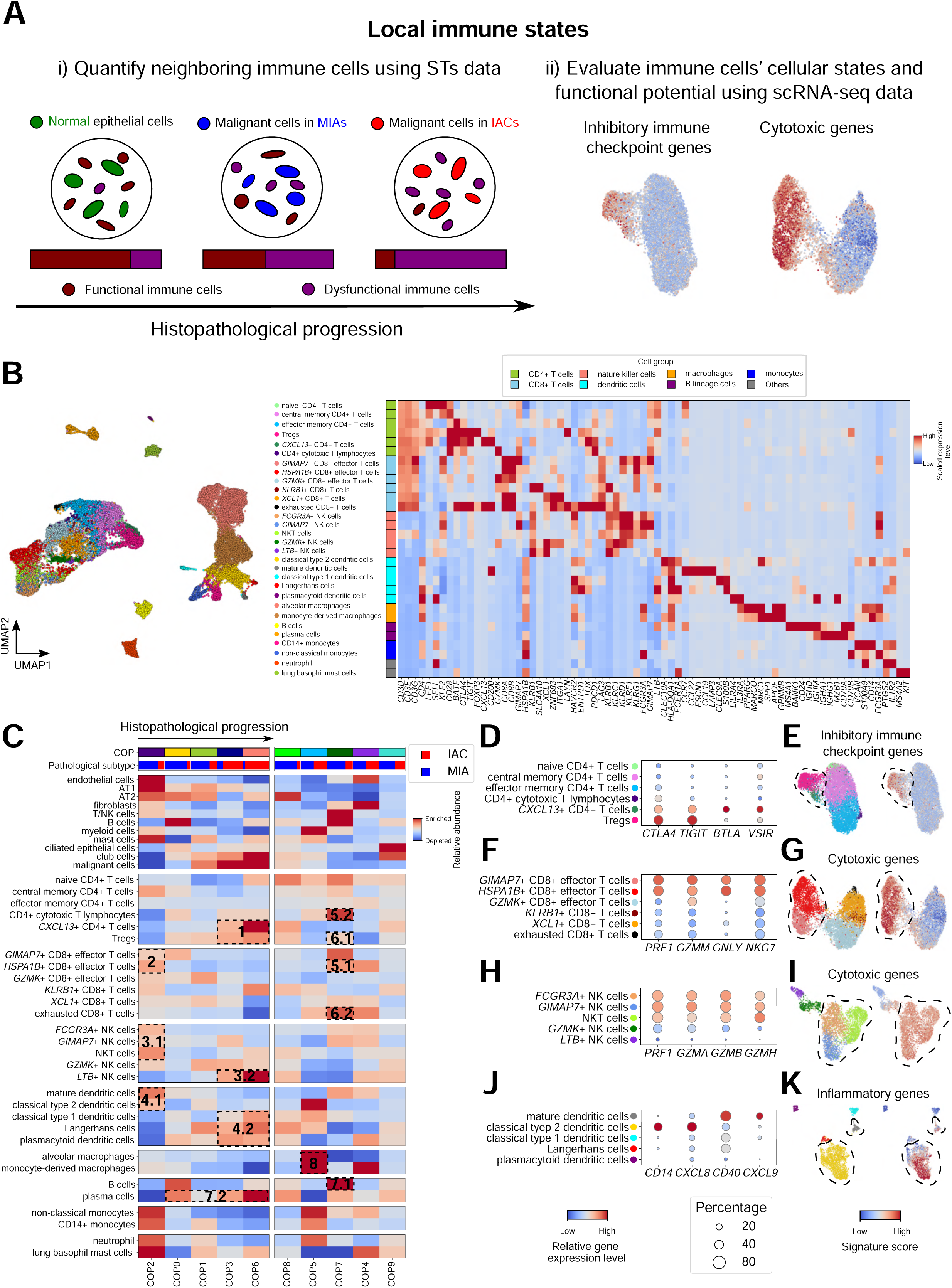
Local protumor immune remodeling during LUAD pathological progression. **(A)** Schematic of neighboring immune cells quantification and cellular states and functional potential evaluation. **(B)** UMAP of 34,545 immune cells from 8 tumor tissues and 3 normal tissues colored by annotated cell types (left panel), expression of marker genes for each immune cell type and colored by cell groups (right panel). **(C)** Heatmap of immune cell enrichment/depletion levels in each COP, bar plot showing proportion of spots from MIA (blue) and IAC (red) patients. **(D)** Bubble heatmap showing the percentage of cells expressing immune inhibitory checkpoint genes (indicated by the size of circle) as well as their relative expression levels (indicated by the color of circle) in each CD4 T cell subpopulation. **(E)** UMAP plot visualization of CD4 T cells colored by annotated cell types and inhibitory immune checkpoint gene scores. **(F)** Bubble heatmap showing the percentage of cells expressing cytotoxic genes as well as their relative expression levels in each CD8 T cell subpopulation. **(G)** UMAP plot visualization of CD8 T cells colored by annotated cell types and cytotoxic gene scores. **(H)** Bubble heatmap showing the percentage of cells expressing cytotoxic genes as well as their relative expression levels in each NK cell subpopulation. **(I)** UMAP plot visualization of NK cells colored by annotated cell types and cytotoxic gene scores. **(J)** Bubble heatmap showing the percentage of cells expressing inflammatory genes as well as their relative expression levels in each dendritic cell subpopulation. **(K)** UMAP plot visualization of dendritic cells colored by annotated cell types and inflammatory gene scores.

### Characterizing the immune neighborhood of malignant/normal epithelial cells and revealing local immunosuppressive remodeling during LUAD progression

The spatial localization of immune cells within TME, including those in close proximity to malignant cells and well-organized immune structures within tumors, is a critical factor for tumor immunity and response to immune checkpoint inhibitors (ICIs)^14,22^. Understanding the spatial organization of immune cells within the tumor microenvironment can provide important insights into the mechanisms underlying tumor immune evasion and resistance to immunotherapy. To address this issue, we integrated the immune cell reference with STs data obtained from patients at different pathological stages of LUAD. Our aim is to delineate the spatial distribution of immune cells within tumor microenvironment and identify the immune states that are in close proximity to malignant cells during cancer progression.

To achieve this, we employed *SPOTlight*^16^ to identify cell type specific topics from scRNA-seq data. These topics were highly specific to each immune cell type, resulting in high specificity for localizing immune cell types in each spot (Supplementary Figure 9A). Then, we examined enrichment or depletion of these immune cells within COPs to understand the local immune states in these regions (Supplementary Figure 9B-C; Details see Methods). We found that the enrichment level of Tregs and *CXCL13*+ CD4+ T cells progressively increased across malignant cells enriched COPs in MIAs (COP0 and COP1), and IACs (COP3 and COP6) compared with normal epithelial cells enriched COPs (COP2) (Figure 2C, box1). These two immune cell types exhibited high expression levels of inhibitory immune checkpoint genes, including *CTLA4*, *TIGIT*, *BTLA* and *VSIR* and showed elevated levels of immune checkpoint gene signature scores (Figure 2D-E). These findings suggest that Tregs and *CXCL13*+ CD4+ T cells may play an important role in establishing a local immunosuppressive microenvironment in LUAD during pathological progression.

Additionally, *GIMAP7*+ CD8+ effector T cells and *HSP1B1*+ CD8+ effector T cells were mainly located in normal epithelial cells enriched COPs (COP2) (Figure 2C, box2). Interestingly, we observed an increase in cytotoxic activity in these CD8+ T cells, as evidenced by higher level of *PFR1* and *NKG7* expression, as well as cytotoxicity signature scores (Figure 2F-G). We further examined the spatial enrichment of NK cell subpopulations in COPs. We also found *FCGR3A*+ NK cells, *GIMAP7*+ NK cells and NKT cells (Figure 2C, box3.1) were primarily enriched in normal epithelial cells enriched COPs (COP2) and showed a higher level of perforin and granzyme expression and enhanced cytotoxicity activity (Figure 2H-I). In contrast, the enrichment level of *LTB*+ NK cells were markedly increased in malignant cells enriched COPs in IACs (COP3 and COP6) (Figure 2C, box3.2). We observed a reduction of cytotoxicity scores in this subpopulation, indicating a decreased ability to eliminate cancer cells, which could contribute the progression of LUAD (Figure 2I).

Our findings also revealed a noticeable variation in the enrichment of different subsets of dendritic cells between normal epithelial cells enriched COPs (COP2) and malignant cells enriched COPs (COP0, COP2, COP3 and COP6) (Figure 2C, box4.[1-2]), suggesting these dendritic cells may have distinct functions in cancer progression. The classical type 2 dendritic cells and mature dendritic cells were found to be gradually depleted from normal epithelial cells enriched COPs (COP2) to malignant cells enriched COPs in MIAs (COP0 and COP1), and last malignant cells enriched COPs in IACs (COP3 and COP6) (Figure 2C, box4.1), while plasmacytoid dendritic cells, classical type 1 dendritic cells and Langerhans cells were observed to be gradually enriched with pathological progression (Figure 2C, box4.2). Increased expression of proinflammatory genes and increased level of inflammatory signature score were evidenced among classical type 2 dendritic cells and mature dendritic cells compared to the other three dendritic cell subsets (Figure 2J-K). These results indicate that as the pathological progression occurs, there is a loss of inflammatory phenotypes in dendritic cells that are in close proximity to malignant cells.

Altogether, these data highlight the presence of local protumor remodeling and immunosuppression during pathological progression, characterized by an increase of immune inhibitory factors of CD4+ T cells and a decrease in cytotoxicity by CD8+ T cells and NK cells as well as a reduction in inflammatory activity by dendritic cells.

We further evaluated the abundance of these immune cells in bulk tissues from different pathological stages using scRNA-seq data. Our observations showed a significant enrichment of CD4+ T cells (i.e., Tregs and *CXCL13*+ CD4+ T cells) expressing inhibitory immune-checkpoints in bulk tumor tissues from both MIAs and IACs, while the proportions of highly cytotoxic CD8+ T cells (i.e., *GIMAP7*+ CD8+ effector T cells, *HSP1B1*+ CD8+ effector T cells) and NK cells (i.e., *FCGR3A*+ NK cells, *GIMAP7*+ NK cells and NKT cells) were decreased (Supplementary Figure 10A-C). Moreover, we noted increased abundance levels of low inflammatory dendritic cells (i.e., plasmacytoid dendritic cells, classical type 1 dendritic cells and Langerhans cells) in LUAD bulk tumor tissues (Supplementary Figure 10D). Importantly, our analysis showed a shift from functional immune cells to dysfunctional immune cells with pathological progression, indicating the presence of immunosuppression components in the LUAD TIME (Supplementary Figure 10E).

### TLSs&LAs signatures are associated with positive response to immune therapy and an active B cell immune response in LUAD

TLSs is an ectopic lymphoid structure that provides an area for the activation and differentiation of T and B cells, which are associated with antitumor immune reactions and responses to immunotherapy^14,15^. In our analysis, we noticed COP7, which was enriched in TLSs and LAs, contained a mixture of both functional and dysfunctional immune cell types (Figure 3A). For example, *HSP1B1*+ CD8+ effector T cells was enriched in COP7 (TLSs and LAs) (Figure 2C, box5.1), which is consistent with a recent report that T cells with high expression of *HSPA1A* and *HSPA1B* are mainly located in LAs across various cancer types, particularly within tumor beds or surrounding tumor edges^23^. However, in our study, these immune cells exhibited high cytotoxicity. Additionally, CD4+ cytotoxic T lymphocytes were also enriched in COP7 (TLSs and LAs) (Figure 2C, box5.2), and this cell type showed high expression of cytotoxicity genes compared to other CD4+ T cells (Supplementary Figure 11). Those immune cell types related to immunosuppression, such as Tregs and exhausted CD8+ T cells, were also enriched in these areas (Figure 2C, box6.[1-2]). These results suggest TLSs and LAs may be a key area for immune reactions in LUAD, where both antitumor responses and immunosuppressive processes coexist. Therefore, further evaluation on their predictive and prognostic value in LUAD progression and immunotherapy is needed.

**Figure 3.**
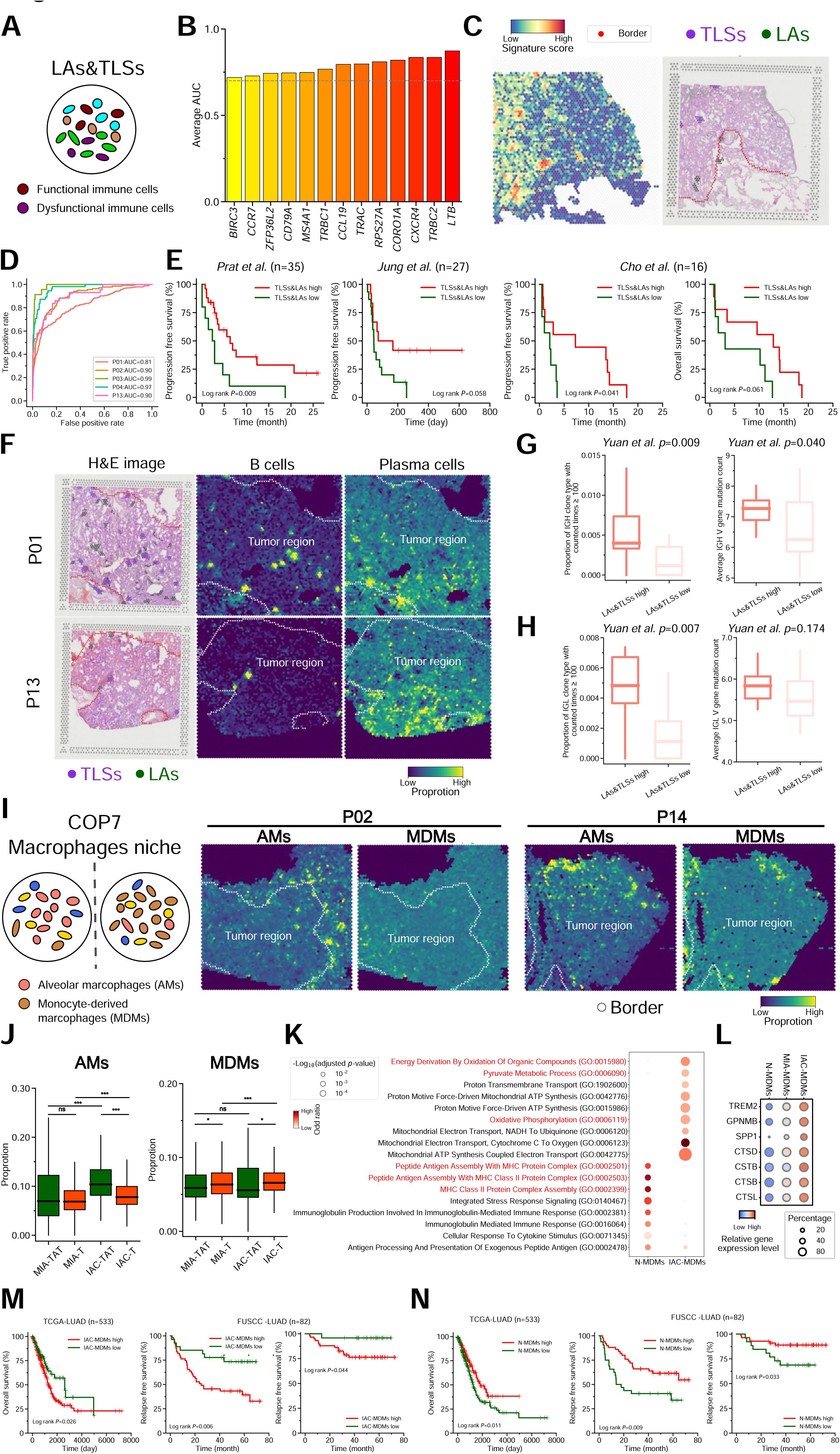
Spatial organization structures are associated with pathological progression and distinct clinical outcomes in LUAD. **(A)** A schematic illustration of colocalization of functional and dysfunctional immune cells in LAs&TLSs (COP7). **(B)** Mean AUC of TLSs&LAs signatures identified from leave-one-out cross validation analyses on TLSs&LAs+ STs data. **(C)** Spatial heatmap of TLAs&LAs signature score and HE image of pathologically identified TLSs&LAs areas in a representative slide (P04). **(D)** ROC curves of TLSs&LAs detection based on the identified signature genes in each slide. **(E)** Kaplan-Meier curves displaying differences in progression-free survival (PFS) or overall survival (OS) between patients whose pretreatment tumors had high or low level of TLSs&LAs signatures scores in the *Prat et al* cohort, the *Jung et al* cohort and the *Cho et al* cohort receiving immune checkpoint inhibitory therapy. **(F)** HE images of pathologically identified TLSs&LAs areas and spatial distribution and proportion of B cells and plasma cells in two representative slides. **(G-H)** Box plots of the proportion of IGH/L clone types counted time greater than 100 (indicate colon expansion) and the average IGH/L V genes mutation counts (indicate somatic hypermutation) between TLSs&LAs high and low samples from *Yuan et al* cohort, *p*-values were derived from rank sum test. **(I)** A schematic illustration of distinct spatial inches for aveolar macrophages (AMs) and monocyte-derived macrophages (MDMs) in COP5. Spatial distribution and proportion of AMs and MDMs in two representative slides. **(J)** Box plots showing the proportions of AMs and MDMs in COP5 spots across tumor (T) and tumor adjacent tissues (TAT) from MIA and IAC patients. **(K)** Functional annotation of differentially expressed genes between MDMs from normal lung tissues (N-MDMs) and IAC tumor tissues (IAC-MDMs). **(L)** Bubble heatmap showing the percentage of cells expressing TAM associated genes and cathepsins as well as their relative expression levels in MDMs from different tissue types. **(M-N)** Kaplan-Meier curves of PFS and OS for TCGA-LUAD cohort and FUSCC-LUAD cohort based on low and high IAC-MDMs (M) or N-MDMs (N) signature scores.

To determine the signature genes of TLSs and LAs, we performed differential gene expression analysis in TLSs&LAs+ samples by leave-one-out cross validation. In brief, we first selected four TLSs&LAs+ samples, and performed differential gene expression analysis between pathologist hand labeled TLSs/LAs spots and other spots in each slide. Only common marker genes were used for further validation analysis. Then we validated these marker genes in the remaining sample and choose those genes associated with an average AUC greater than 0.70 in at least 4 TLSs&LAs+ samples (Supplementary Figure 12A), leading to 13 TLSs&LAs signature genes, including *LTB*, *TRBC2*, *CXCR4*, *CORO1A*, *RPS27A*, *TRAC*, *CCL19*, *TRBC1*, *MS4A1*, *CD79A*, *ZFP36L2*, *CCR7* and *BIRC3* (Figure 3B-C and Supplementary Table 6; Details see Methods). These signature genes were further validated on 5 TLSs&LAs+ samples resulting in AUCs ranging from 0.81 to 0.99 (Figure 3D). To further access the clinical significance of these signature genes, we assessed the association of TLSs&LAs signature with patient response to immunotherapy by analyzing public RNA-seq data sets from four previous clinical studies^24–27^. We found that NSCLC patients exhibited relatively higher TLSs&LAs signature in pretreatment tumors were associated with better response to anti-PD-1 therapy, including significantly better progression-free survival (PFS), overall survival (OS) and durable clinical benefits (Figure 3E and Supplementary Figure 12B).

Previous analysis results revealed that B cells were significantly more abundant in COP7 (TLSs&LAs) (Figure 2C, box7.1). Furthermore, in TLSs&LAs+ samples, B cells were found to be preferentially located at spots annotated as TLSs or LAs, while plasma cells were more diffusely distributed throughout the tumor bed (Figure 3F and Figure 2C, box7.[1-2]). Moreover, when comparing TLSs&LAs+ samples to TLSs&LAs-samples, it was found that B cells in TLSs&LAs+ samples tended to aggregate in LAs and TLSs areas, suggesting an association between TLSs&LAs and B cell immune response (Supplementary Figure 13). The dominance of B cell marker genes (e.g., *MS4A1* and *CD79A*) in the TLSs&LAs signature, as well as enrichment of B cells in LAs and TLSs areas prompted us to focus on B cells immune response. During B cell immunity generation, B cells undergo mutations in their B cell receptor (BCR) sequence, and the most effective clones are selected and expanded^28^. We utilized *MiXCR*^29^ to identify BCR IGL and IGH clonotypes and their somatic hypermutations in RNA-seq data from 36 LUAD tumors^5^, which were classified as either TLSs&LAs high or TLSs&LAs low based on the median expression activity of TLSs&LAs signature. Our analysis revealed that the repertoire of TLSs&LAs high samples exhibited a higher proportion of clonotypes counted more than 100 times in both IGL and IGH (Figure 3G). Furthermore, the overall somatic hypermutation level was higher in TLSs&LAs high tumors compared to TLSs&LAs low tumors, both for IGL and IGH (Figure 3H). These results showed the presence of clonally expanded B cells and somatically mutated IG genes in TLSs&LAs high tumors, suggesting a more active B cell immune response in these tumors. This increased B cell immune activity may be linked to the better immune therapy responses observed in patients with high TLSs&LAs scores.

### Changes of macrophage state are associated with pathological progression and clinical outcomes

Tumor associated macrophages (TAMs) have diverse functions in cancer progression^30–32^. The use of STs and scRNA-seq could advance our understanding of spatial distributions and molecular diversity of TAMs during the progression of LUAD. In our study, we observed that alveolar macrophages and monocyte-derived macrophages were highly enriched COP5 (Figure 2C box8). This finding suggests that these macrophages exhibited a close aggregation, resembling a niche-like local tissue environment (Figure 3I). We conducted a further analysis to determine the density of these two macrophage populations in spots located within the T regions and TAT regions. Our findings revealed that AMs were more highly enriched in spots from TAT regions, particularly in patients with IAC. In contrast, MDMs exhibited a higher enrichment in T spots from both patients with MIA and IAC (Figure 3J). These results suggest that MDMs may serve as important TAMs that were primarily found in the tumor regions. Consequently, we decided to focus our attention on MDMs to gain a deeper understanding of their role in the pathological progression of LUAD. To this end, we performed an analysis to identify DEGs between MDMs derived from IAC patients’ tumor tissues (IAC-MDMs) and normal lung tissues (N-MDMs). As a result, we identified 503 upregulated genes in IAC-MDMs and 253 upregulated genes in N-MDMs (Supplementary Figure 14A and Supplementary Table 7). Notably, the upregulated genes in IAC-MDMs were found to be highly enriched in mitochondrial energic metabolisms, indicating an energy metabolic reprograming in these macrophages during cancer progression, while the up-regulated genes in N-MDMs were primarily associated with MHC-II antigen processing and presentation (Figure 3K). In addition, the expression activity of MHC-II molecules of MDMs decreased with LUAD progression, suggesting a potential impairment in antigen presentation along with the progression (Supplementary Figure 14B). In contrast, the activity of oxidative phosphorylation was up-regulated during pathological progression (Supplementary Figure 14C). MDMs in IAC tended to highly express genes associated with M2 polarization, which is typically associated with an immunosuppressive and protumoral phenotype (Supplementary Figure 14D). More importantly, the expression activity of TAM associated genes (e.g., *TREM2*, *GPNMB* and *SPP1*)^33,34^ and cathepsins (e.g., *CTSD*, *CSTB*, *CTSB* and *CTSL*)^35,36^ were gradually up-regulated along with pathological progression, which have been reported to play an important role in immune suppression in TME (Figure 3L). All these results suggest that MDMs undergo phenotypic and molecular transitions during pathological progression and ultimately transition into an immunosuppressive and protumoral phenotype.

We further evaluate the association of transitional signature with prognosis and found that high expression of the IAC-MDMs signature scores were significantly correlated with worse clinical outcomes in both LUAD-TCGA dataset^37^ and FUSCC-LUAD dataset^38^ (Figure 3M). However, we did not observe a similar correlation with the signature scores of N-MDMs (Figure 3N). Altogether, these results indicate that MDMs exhibit unique spatial characteristics and molecular diversity during LUAD progression. In addition, the energy metabolism of MDMs may be a potential immunotherapy target to reverse M2 polarization and immune suppression in TIME.

### Cell-to-cell cross talk during LUAD pathological progression

We further explored the potential mechanisms underlying the differential spatial distribution and functional properties of immune cell subpopulations in COPs. These regional immune cell distributions may relate to local immunoregulation. We first focus on the molecular alterations of epithelial cells during pathological progression, and signature genes of MIA and IAC malignant cells were identified. These signature genes were then used to estimate the expression activity in bulk RNA-seq samples obtained from a previous study^38^, which included samples from adenocarcinoma in situ (AIS), MIA and IAC. Consistently we observed higher MIA malignant cell signature scores in AIS/MIA samples compared to IAC samples. Conversely, IAC malignant cell signature genes were highly expressed in IAC samples in this independent data set (Figure 4A). Furthermore, we also evaluated the correlation between the signature scores and functional molecular markers of immune cell. Our analysis revealed a more significant positive correlation between IAC malignant cell signature scores and the expression of *FOXP3* which is associated with immune suppression (Figure 4B). Conversely, we observed a stronger negative correlation between the IAC malignant cell signature scores and the expression of antitumor immune markers (*PRF1* and *GZMA*), indicating a potential decrease in cytotoxicity (Figure 4C). These results suggest that the potential impact of these molecular characteristics of IAC malignant cells on the immune suppression within the TIME.

**Figure 4.**
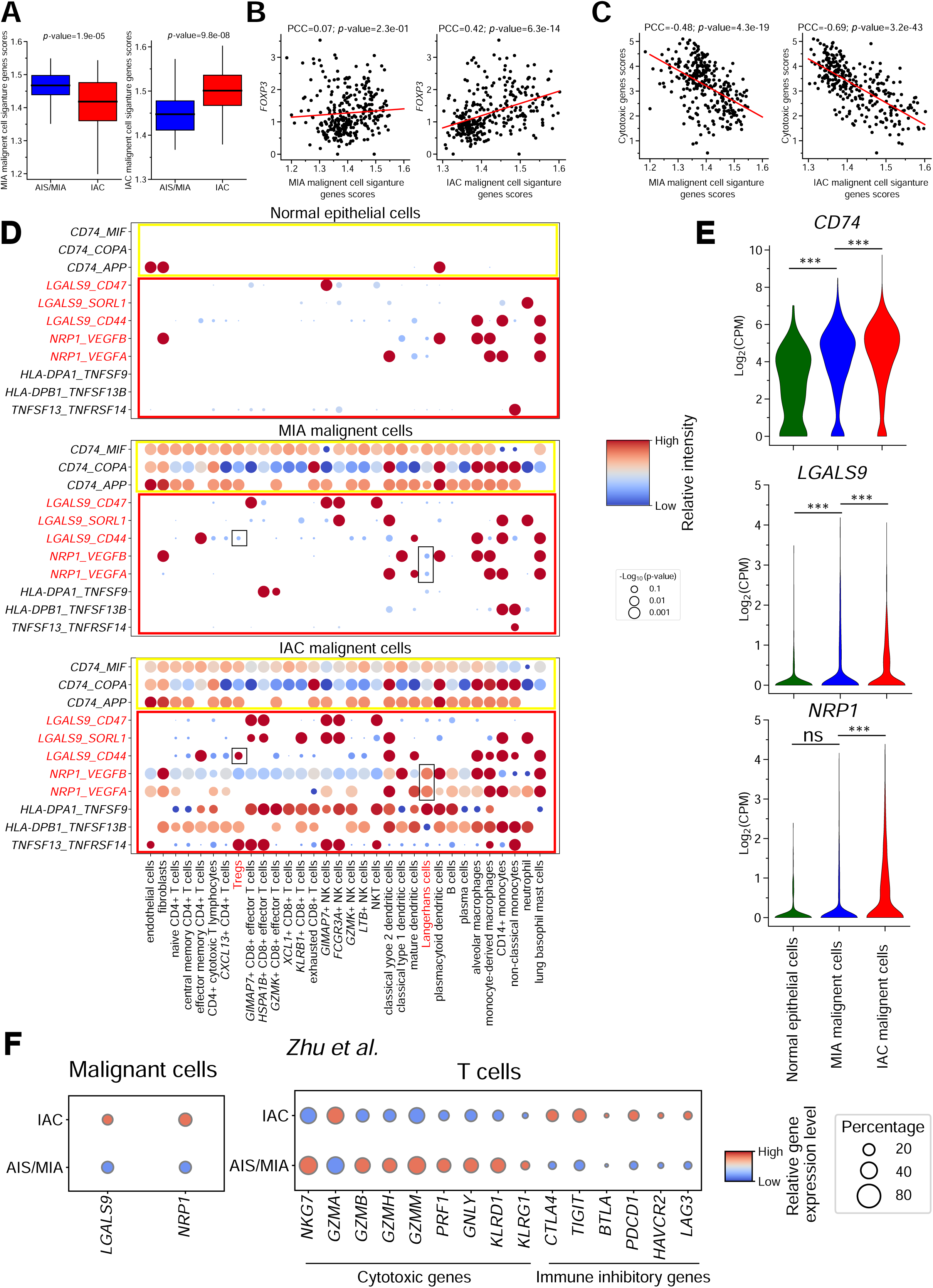
The alterations in malignant cells molecular characteristic and changes in crosstalk between malignant cells and TME cells during LUAD progression. **(A)** Boxplots showing signature scores of scRNA-seq data defined MIA/IAC malignant cell marker genes in an independent cohort (FUSCC-LUAD). **(B-C)** Scatter plots of correlation of expression using Pearson correlation coefficients between MIA/IAC malignant cell signature and cytotoxic score (B), *FOXP3* (C) in FUSCC-LUAD cohort. **(D)** Overview of the selected IAC malignant cells upregulated ligand-receptor interactions, *p*-values are indicated by circle size, relative interaction strengths indicated by color. **(E)** Expression levels of three immune modulators in normal epithelial cells, MIA malignant cells and IAC malignant cells. **(F)** Bubble heatmap showing the percentage of cells expressing immune modulators, cytotoxic genes and immune inhibitory genes as well as their relative expression levels in malignant cells and T cells across different progression stages in an independent dataset.

The interactions between malignant cells and elements in TIME can lead to the development of an immunosuppressive phenotype that aids in tumor progression. Thus, we used *CellPhoneDB*^39^ to identify the expression of potential crosstalk signaling molecules in malignant/normal epithelial cells based on ligand-receptor interactions. Notably, our analysis revealed an increase in the interaction between malignant cells and other cell types with tumor progression (Supplementary Figure 15). Next, we focus on those interactions specific to IAC malignant cells. We identified altered interactions that exhibited differential modulation in malignant cells, particularly in IAC. Notably, we observed an enhanced interaction between *CD74* on malignant cells and *MIF*, *COP*, and *APP* on other cell types. Additionally, we discovered increased interactions between immune modulators *LGALS9* and *NRP1* on IAC malignant cells, and *CD44* on Tregs, as well as *VEGFA/B* on Langerhans cells. (Figure 4D). Furthermore, the expression activity of these immune receptors was progressively upregulated in malignant cells across pathological stages (Figure 4E). These immune modulators have been reported to play an important role in lung cancer immune suppression^40,41^. We further validated the expression levels of *LGALS9* and *NRP1* in malignant cells at different progression stages using an independent dataset^10^ (Supplementary Figure 16). Results in this dataset consistently demonstrate increased expression of immune modulators in malignant cells of IAC. Furthermore, we observed a negative correlation between the expression of these immune modulators and cytotoxic gene expression in T cells, while a positive correlation was observed with the expression of immune inhibitory genes (Figure 4F). Altogether, these results suggest that altered cell-to-cell interactions occurs during pathological progression, which may be an important factor in local immune remodeling.

### Phenotypic clusters from heterogeneous malignant cells are associated with pathological progression and clinical outcomes

The presence of interpatient heterogeneity among malignant cells from different patients hindered the ability to directly compare them. To overcome this, we employed an integration method that takes into consideration the individual characteristics of each patient to identify distinct phenotypic clusters within malignant cells^42^. Through this approach, we successfully identified eight phenotypic clusters (referred to as PCs, PC[0-8]) (Figure 5A). Notably, the proportions of PC1, PC4, and PC6 were found to be higher in IAC (Figure 5B). We further characterized PCs based on hallmark pathway activity. WNT beta catenin signaling, epithelial-to-mesenchymal transition (EMT) and IL6-JAK-STAT3 signaling were highly activated in PC4 cells (Figure 5C), suggesting the potential invasive ability of PC4 cells^43,44^. In order to examine the relationships between phenotypic clusters and clinical outcomes, we first identified a set of differentially expressed genes that served as signature genes for each phenotypic cluster. Subsequently, we assessed the expression activity of these signature genes in individual samples obtained from two publicly available datasets^37,38^. Our analysis revealed a consistent and significant negative correlation between PC4 signature genes and both PFS and OS in patients in two independent datasets (Figure 5D).

**Figure 5.**
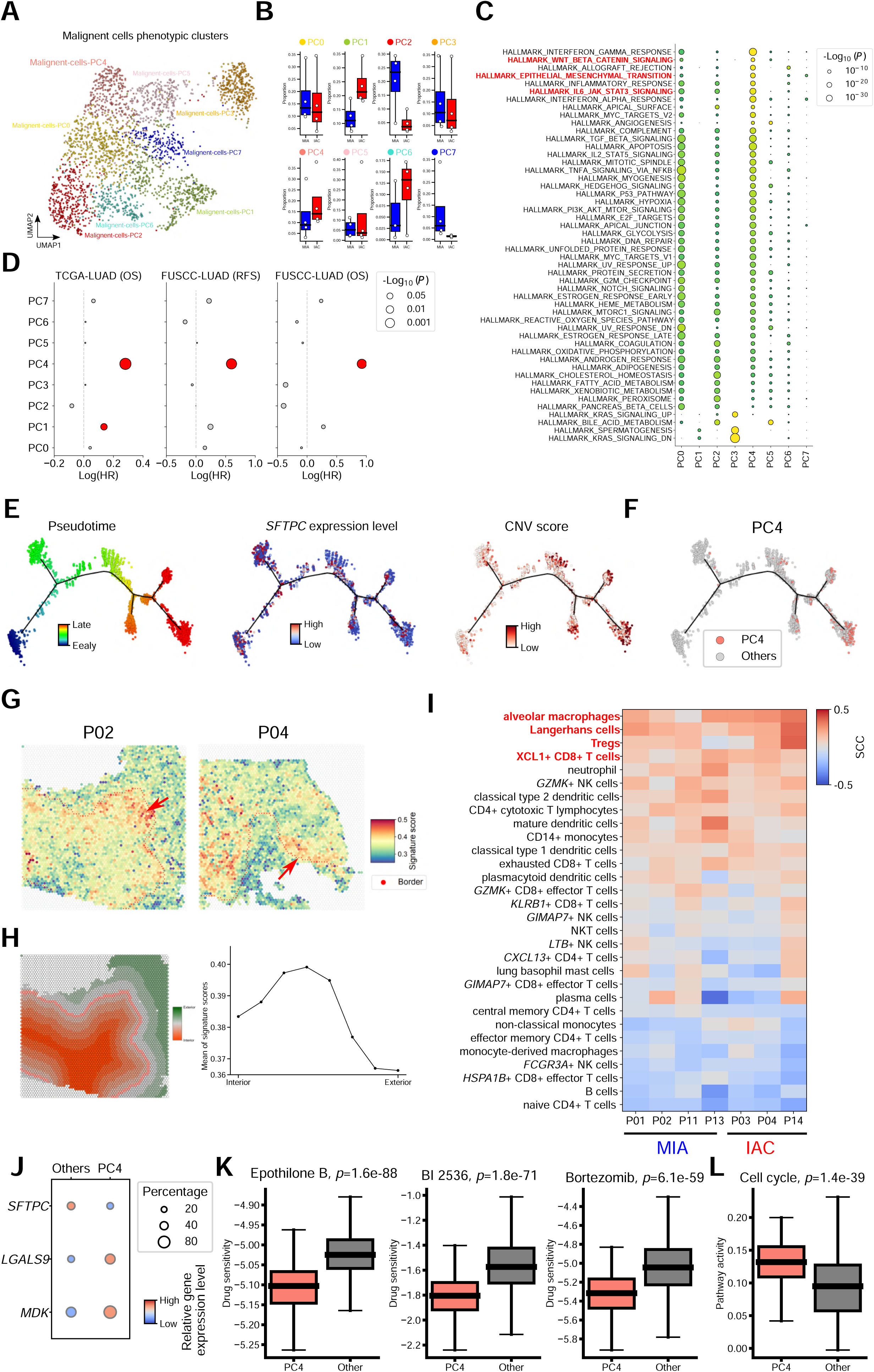
Integration method identified a unique leading edge malignant cell state during LUAD pathological progression. **(A)** UMAP of malignant cells after removing patient specific signals by integration method. Cells are colored by identified clusters (phenotypic clusters). **(B)** The proportion of eight malignant cells phenotypic clusters in MIA and IAC patients. **(C)** Heatmap of hallmark pathway activity among phenotypic clusters. **(D)** Hazard ratios of all phenotypic cluster signatures in two LUAD cohorts. **(E)** Trajectory analysis of malignant cell phenotypic clusters, cells were colored by their pseudotime, SFTPC gene expression level and CNV score. **(F)** Trajectory plot showing the distribution of PC4 on the malignant cell progression trace. **(G)** Spatial heatmaps of PC4 signature scores in two representative slides. Red dots indicate the border of tumor region (leading edge). **(H)** Dividing spots into eight groups based on their distances to the tumor boundary and the mean of PC4 signature scores in each group. **(I)** Correlations between PC4 signature scores and the proportions of other 30 immune cell type. **(J)** A boxplot showing the predicted drug sensitivity score (IC50) for PC4 and other malignant cells based on their gene expression profiles. **(K)** A boxplot showing the cell cycle pathway activity of PC4 and other malignant cells based on their gene expression profiles, *p*-values were derived from rank sum test in both (J) and (K).

In order to further investigate the key molecular event in the progression of malignant cells, we performed pseudotime trajectory analysis on all malignant cell clusters using Monocle2^45^ and revealed a gradually loss of AT2 marker expression and increase of CNV scores along the pseudotime (Figure 5E). Cells from PC4 were gathered on the end of trajectory and associated with highest CNV scores among all malignant cell phenotypic clusters (Figure 5F and Supplementary Figure 17A). Specifically, the CNV scores of PC4 were further increased in IAC (Supplementary Figure 17B). These results suggest that PC4 might be a subpopulation progressively accumulates with LUAD progression and helps drive LUAD invasion.

Further spatial distribution analyses found that PC4 signature genes were highly enriched in the border between tumor regions and adjacent normal regions. Besides, peripheral tumor regions also expressed relatively high levels of this signature (Figure 5G-H). These results suggest that PC4 may be a population aids malignant cell invasive at leading edge of tumor tissues. To further evaluate the local immune state of PC4 enriched regions, we correlated the signature score with immune cell abundance. Interestingly, alveolar macrophages, Langerhans cells and Tregs were the top-ranked correlated immune cells, indicating a local immune suppressive state (Figure 5I). Alveolar macrophage was the most positively correlated immune cell type, and the correlations were higher in IAC patients than those in MIA. These results were in line with a recent report that alveolar macrophages accumulate close to tumor cells early during tumor formation to promote epithelial–mesenchymal transition and invasiveness in tumor cells, also induce a potent suppressive Treg response^21^. We also found the expression level of immune suppressive modulators *MDK* and *LGALS9* were upregulated in PC4, indicating an immune suppression phenotype (Figure 5J)^46–48^. Spatial analysis of immune cell types and states of leading edge of tumor regions reveals a local suppressive immune environment involved in promoting cancer cell invasion.

In order to identify which drugs might be useful to suppress PC4 malignant cells, *pRRophetic*^49^ was applied to predict drug sensitivity from malignant cell gene expression levels. Our analysis showed that *Epothilone B*^50^, *BI 2536*^51^ and *Bortezomib*^52^ had significantly lower IC50 values in PC4 than that in other PCs (Figure 5K), suggesting that PC4 has the higher sensitivity to these three drugs. These three drugs have been reported to prevent cancer cells from proliferating by interfering with cell cycle. Consistently, the cell cycle activity was significantly upregulated in PC4 cells (Figure 5I). Considering the presence of a localized suppressive TIME remodeling and an accumulation of invasive malignant cell population in the progression of LUAD, the combination of chemotherapy and immunotherapy holds promising potential as an effective therapeutic approach for advanced or metastatic cases of LUAD (e.g., IAC).

## Discussion

The understanding of local immune remodeling in LUAD pathological progression is limited. In this study, we presented a comprehensive spatial cellular organization map in MIA and IAC by integrating STs data and scRNA-seq data of tumor tissues and normal lung tissues from eight LUAD patients. In total, we identified ten COPs that recapitulated both known and new tissue architectures, including well-organized immune cell structures, such as LAs and TLSs. By comparing the immune states and functional potential of those immune cells enriched or depleted in malignant cell enriched COPs, we observed the presence of local protumor remodeling and immunosuppression during pathological progression that comprised the increase of immune inhibitory factors of CD4+ T cells and the decrease in cytotoxicity by CD8+ T cells and NK cells as well as a reduction in inflammatory activity by dendritic cells (Figure 6I). Thus, investigating the local immunosuppressive remodeling during LUAD pathological progression may facilitate the discovery of protumor immune components and targets for immune therapy.

**Figure 6.**
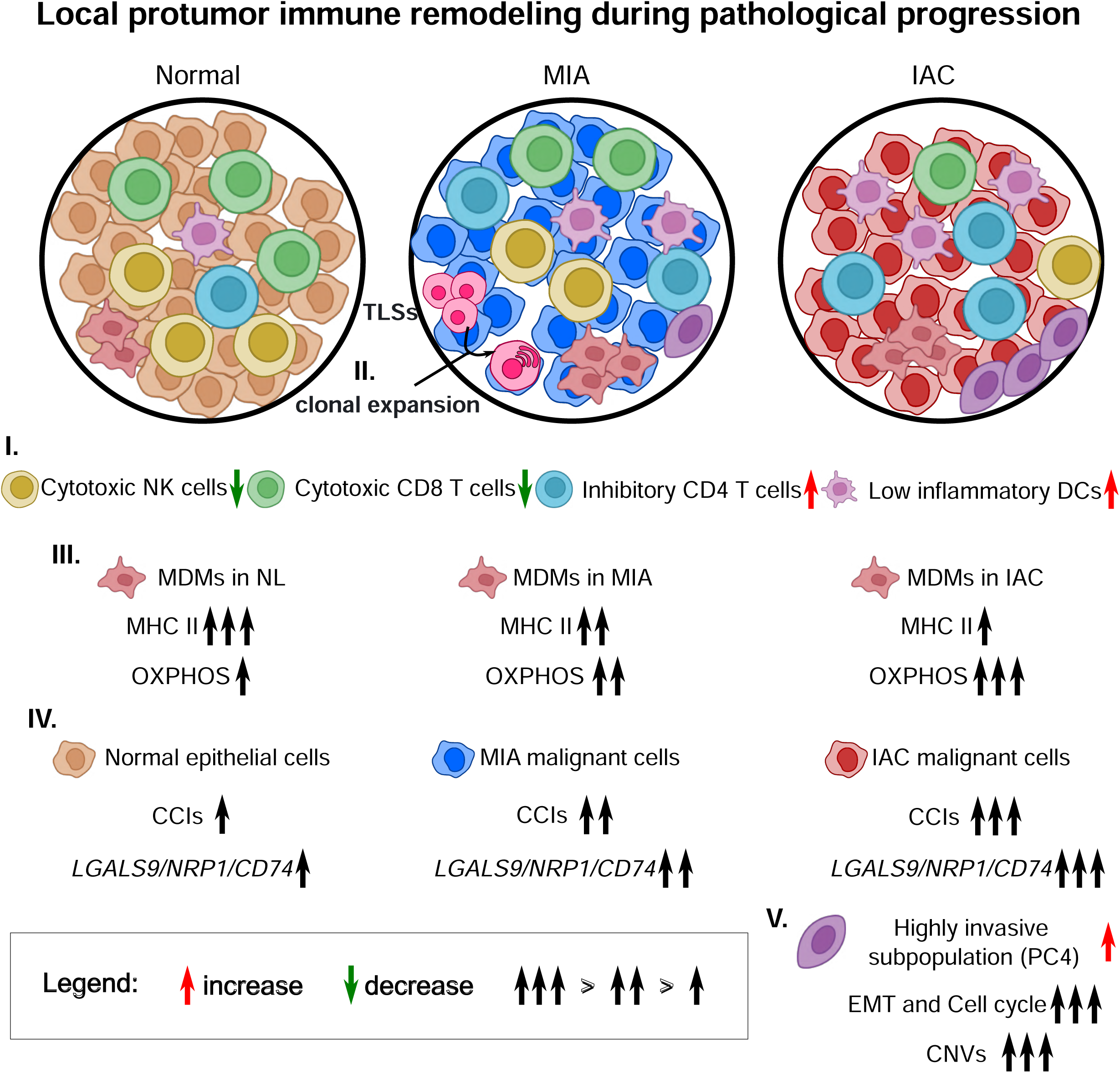
Schematic overview of spatial cellular organization and functional states transition involved in the cancer progression of LUAD. Local pro-tumor immune remodeling during pathological progression. **I**. Depletion of functional immune cells (Cytotoxic NK cells and CD8 T cells) and enrichment of dysfunctional immune cells (Inhibitory CD4 T cells and Low inflammatory DCs) surrounding malignant cells with LUAD progression. **II**. Association between TLSs and B cell clonal expansion. **III**. The changes of macrophage state during LUAD progression, OXPHOS, oxidative phosphorylation. **IV**. Increased number of CCIs and expression of immune modulators in malignant cells during LUAD progression, CCIs, cell-to-cell interactions. **V**. A specialized malignant cell phenotypic cluster located at the boundary of tumor tissues, EMT, epithelial-mesenchymal transition, CNVs, copy number variations.

It has been reported that high-order arrangements of cells, such as lymphocyte aggregates (LAs) and tertiary lymphoid structures (TLSs), represent structural and functional components in tumor tissues and contribute to a positive immune therapeutic response and clinical outcomes^14,15,53,54^. In this study, we identified COP7, which was highly colocalized with TLSs and LAs, and was associated with both functional and dysfunctional immune cell types. However, TLAs&LAs signature defined by STs were positively correlated with immune therapy outcomes in several independent ICI data sets. Notably, B cells were markedly enriched in TLSs&LAs regions, especially in TLSs, while plasma cells were diffusely distributed throughout the tumor bed, suggesting a potential activated B cell immune response in TLSs&LAs+ samples. In the present work, we validated the a more active B cell immune state in TLAs&LAs-high LUAD samples, including the presence of clonally expanded B cells and somatic hypermutated BCRs (Figure 6II). It is also accordance with several previous studies showing that infiltration of B cells and plasma cells correlated with TLSs presence^14,53^. Altogether, B cell immune responses contribute to antitumor function in TLSs presence tumor samples and promotes immune therapy efficiency in these samples.

Tumor-associated macrophages (TAMs) play diverse roles in cancer progression, with their functions and states being influenced by their specific locations ^21,30^. In this study, we identified two distinct macrophage populations: alveolar macrophages (AMs), which are tissue resident macrophages (TRMs) and monocyte-derived macrophages (MDMs). We found both AMs and MDMs exhibited high enrichment of COP5, indicating their close aggregation and resemblance to a niche-like local tissue environment, representing a potential therapeutic target. MDMs were associated with energy metabolism (e.g., oxidative phosphorylation, OXPHOS) changes as disease progressed, and there was a reduction in the expression of MHC II molecules as disease progressed, suggesting a potential impairment in the antigen presenting process (APC) (Figure 6III). Alterations in OXPHOS and APC might skew their function toward a protumor phenotype. Consistently, higher expression activity of IAC-MDMs signature correlated with shorter OS and PFS in two independent datasets. The energy metabolism of MDMs may be a potential immunotherapy target to reverse M2 polarization and immune suppression in TIME.

Demonstrating cell-to-cell interactions (CCIs) can provide important insights into tumor-immune coevolution and immune remodeling^7,8^. Our findings indicate a progressive increase in the number of predicted interactions between malignant cells and other cells within TME with pathological progression. Notably, the strength of interactions involving immune checkpoint receptors, such as *LGALS9* and *NRP1*, was found to be heightened in patients with invasive adenocarcinoma (IAC) (Figure 6IV). Furthermore, *LGALS9* expression was significantly elevated in malignant cells of IAC patients. It is worth noting that *LGALS9* has been reported to be associated with immunomodulatory activity and cancer immune evasion^40^. These results suggest potential therapeutic targets for immune-based treatments that could enhance the development of an anti-tumoral TIME.

The presence of interpatient heterogeneity among malignant cells from different patients posed a challenge in directly comparing them. To addressed this issue, we implemented an integration method that takes into consideration the individual characteristics of each patient to identify distinct phenotypic clusters within the malignant cells. Utilizing this strategy, we discovered a subgroup of malignant cells (i.e., PC4) located at the boundary of tumor tissues, which exhibited enhanced invasive potential (Figure 6V). Notably, the expression activity of the signature genes associated with this phenotypic cluster showed a significant negative correlation with OS and PFS in other independent datasets. Furthermore, these cells were found to colocalize with AMs. This finding aligns with a recent study highlighting the accumulation of tissue resident macrophages (TRMs) in close proximity to tumor cells during early tumor formation, promoting EMT and tumor cell invasiveness and inducing a Treg response. Furthermore, we found that PC4 cells showed increased sensitivity to drugs that target the cell cycle pathway. This suggests that combining chemotherapy and immunotherapy may hold promising potential as an effective therapeutic approach for advanced or metastatic cases of LUAD, particularly those with an IAC subtype.

In this study, although the 10X Visium technology used did not achieve single cell resolution, spatial cellular organization is the crucial factor that determines tumoral immune response and cancer invasive progression. To overcome this limitation, we employed a deconvolution strategy to maximize the resolution and provide the spatial location and composition of different cell types within tumor tissues. Moreover, due to the inherent variability in sample collection, it was challenging to robustly demonstrate the progression trajectory using a small number of patient samples. However, we were able to integrate omics data from multiple tumor tissues at different pathological stages, which help us to dissect the recurrent spatial cellular organization in LUAD and gain valuable insights into the progression of the disease. At last, our findings could be further validated in animal models, which provide an opportunity to experimentally manipulate and investigate the spatial cellular organization in LUAD, allowing for a more comprehensive understanding of the disease progression and potential therapeutic interventions.

To identify potential therapeutic targets, it is essential to comprehend the dynamic spatial cellular organization of cells within TME. In our study, we utilized a combination of STs data and scRNA-seq data to gain insights into specialized cells within LUAD TME. This integration allowed us to understand how these cells are organized in space, how their functional states vary across LUAD progression and how these structures impact clinical outcomes. By employing this approach, we have achieved a deeper understanding of the structural aspects of immunity in LUAD, which in turn provides a valuable resource for the identification of potential therapeutic targets.

## Methods

### Patient enrollment and sample collection

Eight patients who underwent surgery in the Department of Thoracic Surgery of Fudan University Shanghai Cancer Center were enrolled in this study. The inclusion criteria were as fallowing: (1) mixed ground-glass opacity (GGO) featured lung nodule; (2) treatment naïve; (3) maximum diameter of the tumor ≤ 3.0 cm; (4) the frozen section was diagnosed as minimally invasive adenocarcinoma (MIA) or invasive adenocarcinoma (IAC), and verified with paraffin section by pathologists.

This study was reviewed and approved by the Institutional Review Board of FUSCC. Each patient has signed the informed consent form before tissue collection for research studies. Freshly tumor (n = 8) and adjacent normal lung samples (n = 8) were delivered within MACS^®^ Tissue Storage Solution (Miltenyi Biotec) on ice directly from the operating room to the laboratory after frozen pathology (1-2h).

### Sample handling

Each obtained tumor sample was divided into three pieces and processed for spatial transcriptomics (ST), single-cell RNA sequencing (scRNA-seq), and bulk sequencing, respectively. The following steps are performed on ice. Briefly, tumor pieces for scRNA-seq were rinsed with cold phosphate-buffered saline (PBS), minced into pieces (approximately 1 mm3) and set aside in PBS for preparing single-cell suspensions. Tumor pieces for ST were embedded in optimal cutting temperature compound (OCT) compound (SAKURA) and then snap-frozen on dry ice immediately and stored at -80°C until cryosectioning. The remaining tumors were frozen at -80°C for bulk sequencing (RNAseq and WES).

Meanwhile, all paired adjacent normal lung samples were processed for bulk sequencing, among which three samples (P02, P04 and P12) were processed for scRNA-seq as mentioned above.

### Single-cell suspensions for scRNA-seq

1 mm3 tumor pieces were enzymatically digested with 20ml digestion medium containing 250 U/ml collagenase I, 100 U/ml collagenase IV (Worthington) and 30 U/ml DNase I (Worthington) for 45 min at 37°C in a shaking water bath with agitation. Then, the suspension was filtered through a 70-μm cell strainer, and centrifuged at 300g for 8 min. After removing the supernatant, the cell pellet was suspended in red blood cell lysis buffer (Miltenyi Biotec) for 5 min and centrifuged at 300g for 8 min. the pellet was washed twice and re-suspended with PBS with 0.04% BSA. After re-filtered through a 35μm cell strainer, the dissociated single cells were stained with AO/PI for cell concentration and viability assessment through Countstar Fluorescence Cell Analyzer (ALIT). The cell concentration was adjusted to 1000 cells/μl approximately.

### scRNA-seq

Library construction was performed according to the protocol of 10X Genomics Chromium Controller Instrument and Chromium Single Cell 3’ V3 Reagent Kits (10X Genomics). After quantified by the Qubit High Sensitivity DNA assay (Thermo Fisher Scientific) and Bioanalyzer 2200 (Agilent), the constructed libraries were sequenced on NovaSeq 6000 (Illumina) platform.

### Tissue section preparation, fixation, staining and imaging

Cryosections were cut in the cryostat (Leica) with a thickness of 10 μm. Firstly, RNA integrity number was assessed with 10 slides through the RNeasy Mini Kit (QIAGEN), of which the acceptable quality level is ≥7. Secondly, the permeabilization time was optimized with Visium Spatial Tissue Optimization Reagent Kits (10X Genomics). Sections on Visium Spatial Tissue Optimization Slides (10X Genomics) were fixed, stained, and then permeabilized for different times. The degree of mRNA release is transferred to a fluorescence signal and captured by the fluorescent microscope (ECLIPSE Ti, Nikon). The permeabilization time that produces the highest fluorescence signal with minimum signal diffusion was considered optimal. In this study, 6 min is optimal permeabilization time. Then, tissue sections on Visium Spatial Gene Expression Slides (10X Genomics) were incubated at 37°C 1 min, fixed in methanol at -20°C for 30 min. The slides were stained after being incubated in isopropanol for 1 min, Hematoxylin for 7 min, Bluing Buffer for 2 min, and Eosin for 1 min. After each staining step, slides were washed with DNase and RNase free water. Stained tissue sections are imaged by the microscope (ECLIPSE Ti, Nikon) and carefully reviewed by pathologist to annotate as tumor tissue, tumor adjacent tissue, bronchiole, coal dust, vascular, lymphoid aggregates and tertiary lymphoid structures.

### ST library construction and sequencing

Tissue permeabilization cDNA Synthesis, Library Construction were performed with Visium Spatial Gene Expression Reagent Kits (10X Genomics) and Library Construction Kit (10x Genomics) according to the Visium Spatial Gene Expression User Guide (CG000239, Rev B, 10x Genomics). the constructed libraries were sequenced on NovaSeq 6000 (Illumina) platform.

### Whole exome sequencing and data processing

DNA from tumors and paired adjacent normal lung samples was extracted using TIANamp Genomic DNA Kit (TIANGEN) following the manufacturer’s instructions. Exon library was constructed using NEBNext® Ultra™ II DNA Library Prep Kit for Illumina (NEB). DNA library hybridization was performed by using Agilent SureSelect Human All Exon V6 (Agilent). Paired-end sequencing (150 bp) was performed on the Illumina NovaSeq platform. For WES analysis, sequencing reads were aligned to the hg38 reference genome using *bwa*^55^ (v.0.7.17). *Picard* (v.2.27.4) was utilized for marking putative duplications. Sites that potentially contained small insertions or deletions were realigned and recalibrated using *GATK*^56^ (v.4.3.0) modules. The recalibration process involved the utilization of known variant sites from 1000G phase1 high confidence snps and Mills and 1000G gold standard indels. Somatic mutations were called using *GATK Mutect2*.

### RNA-seq and data processing

Total RNA from tumors and paired adjacent normal lung samples was extracted using TRIzol reagent (Invitrogen). The cDNA libraries were constructed by using the TruSeq Stranded mRNA Library Prep Kit (Illumina) according to the manufacturer’s instructions. The libraries were sequenced on HiSeq X platform (Illumina) and 150 bp paired-end reads were generated. RNA-seq reads were aligned to the reference human genome (hg38) with *STAR*^57^ (v.2.6.1a). Then *featureCounts*^58^ (v.1.6.4) was used for quantification and *DESeq2*^59^ was used for differential analysis.

### Spatial transcriptomics data processing and basic analysis

Raw Visium Spatial RNA-seq output and bright field of images of hematoxylin and eosin (H&E) stained biopsy were processed by *spaceranger* (V1.2.0) to detect tissue, align reads and generate feature-spot matrix. Basic QC was applied on these data. In detail, spots with low transcript content (UMIs count less than 1,000) were removed and HE stained images were labeled by a pathologist to indicate which spots contained cancerous and noncancerous tissues. Then we defined tumor signature genes and TAT signature genes by performing differential expression genes analysis between bulk RNA-seq profiles of bulk tumor and bulk normal lung tissues (Supplementary Figure2). To verify whether the spatial transcriptomic features are consistent with the histological information, we designed a model to distinguish cancerous spots and non-cancerous spots using tumor signature genes score and normal lung signature genes score. We used the Area Under the Curve (AUC) to measure the ability of this model to distinguish between two classes. A randomly selected genes set was used as a negative control.

### Raw single-cell sequencing data processing and basic analysis

Raw scRNA-seq data were preprocessed, demultiplexed, align to human reference genome (hg38) by using *cellranger* (v.4.0.0^11^. Cellular debris and doublets were considered as outliers and outliers were detected using the median absolute deviation (MAD) from the median. For a given metric (e.g., library size or the number of detected genes), outlier was one that lies over two number of MAD away from median. Low quality cells where greater than 10% of transcripts were derived from the mitochondrial genome were excluded. Finally, 45,053 cells were retained for downstream analysis.

Raw UMI counts were normalized and scaled and used for PCA using *Seurat* (v.3.2.3)^60^. Unsupervised clustering analysis were performed. UMAP was used for visualization for clusters. First, we identified DEGs for each cluster using *FindAllMarkers* function in *Seurat* and inferred the major cell types of each cluster based on the top-ranked DEGs. Totally, we defined 8 major cell types, including T/NK cells (*CD3D*/*CD3E*), B lineage cells (*MZB1*/*MS4A1*), mast cells (*TPSB2*/*TPSAB1*), myeloid cells (*CD68*/*S100A8*/*S100A9*), endothelial cells (*VWF*/*PECAM1*/*CD34*), fibroblasts (*DCN*/*COL1A2*/*COL3A1*), ciliated epithelial cells (*TPPP3*/*FOXJ1*) and epithelial cells (*EPCAM*/*SFTPB*).

### Analysis of copy number variations (CNVs) in epithelial cells

Copy number variations (CNVs) in epithelial cells were inferred using *inferCNV* (v.1.2.1)^12^ with endothelial cell as a control. To quantify the level of CNVs, we first computed the mean of squares of relative CNV values across each chromosome (chrX and chrY were excluded in this analysis) in an epithelial cell and then aggregated all chromosomes into an average score.

More specifically, if we let *CNV_g_* denotes the CNV value of gene *g* in epithelial cell *i*, the CNV score of epithelial cell *i* is given in Equation (1):

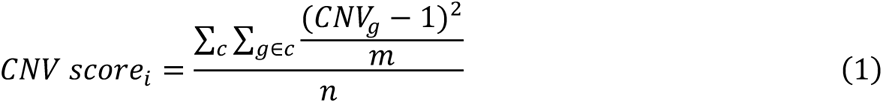

where *m* is the number of genes located at chromosome *c* and *n* is the number of autochromosomes.

### Spatial transcriptomics spots deconvolution using *SPOTlight*

To estimate the cellular composition in each spot in STs data, *SPOTlight*^16^ was used to deconvoluted transcriptome profile in each spot into a combination of cell type specific transcriptome profiles. We first acquired the marker genes of each cell type in scRNA-seq data, then further training the cell type topic profiles and deconvolute the spots into a cellular composition using the function *spotlight_deconvolution*. The same analysis procedure was also applied to estimate the local immune cell composition in each spot.

### Cellular organization patterns identification

All deconvolution results were integrated and clustered into ten cellular organization patterns (COPs) using standard scRNA-seq analysis pipeline in Seurat. Then we created a two-dimensional embedding of spotwise cellular compositions using uniform manifold approximation and projection (UMAP) for visualization. Spots with similar cell type composition were located close to each other in this UMAP space (Supplementary Figure6).

### Linking COPs to pathologically annotated areas

To link the COPs to pathologist-annotated areas, we assessed the degree of enrichment or depletion of annotated areas in each COP by the following approach. Let *N* be all spots present in STs data, let *P* be the set of spots associated with a pathologically annotated area and let *C* be the set of spots in a COP. Then conduct a Fisher’s exact test (two-sided) to evaluate the significance of overlap between *P* and *C*. For each COP construct a 2X2 contingency table is illustrated in Table 1:

**Table 1.** Contingency table used for evaluate the degree of enrichment or depletion of annotated areas in each COP.

|  | Spots in COP | Spots not in COP |
| --- | --- | --- |
| **Associated with P** | $\vert C \cap P\vert $ | $\vert (N - C) \cap P\vert $ |
| **Not Associated with P** | $\vert C \cap (N - P)\vert $ | $\vert (N - C) \cup (N - P)\vert $ |

Odd ratios (*OR*) and p-values (*p*) obtained from above analyses were transformed into enrichment/depletion scores as presented in Equation 2:

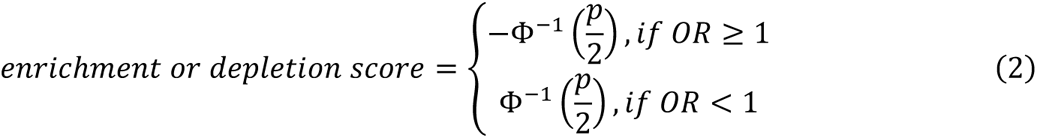

where −Φ^−1^ is the inverse cumulative distribution function of standard normal distribution.

### Quantifying the immune cell infiltration level in each COP

To quantify the infiltrating level of cell types (immune cell types) in each COPs, for a specific cell type, we first calculated the average proportion value as the observed average. Then we permuted the labels of spots and randomly selected a set of spots that matched the number of spots in the COP, computed the average proportion value as the permuted average, repeated this procedure 1,000 times. The differences between observed average and permuted average were computed and the mean value of the differences scaled by the its standard deviation was taken as the relative infiltrating score for each COP. Specifically, for a certain cell type, let *μ_observed_* denotes the average proportion value of this cell type and let *μ^i^_permuted_* denotes the average proportion value in *i^th^* permutation. The relative infiltrating score is defined as follow:

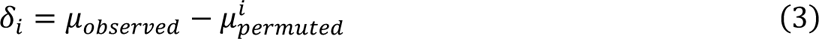

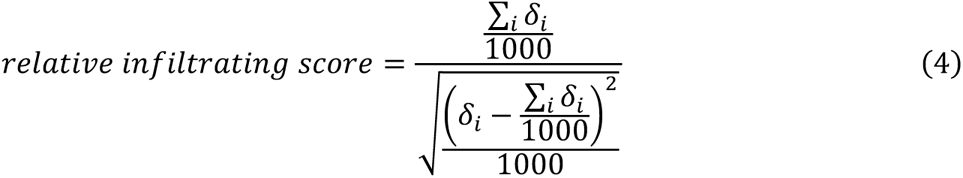

Positive scores indicate relative enrichment but negative scores represent relative depletion (Supplementary Figure 8B-C, Figure 1H and Figure 2C).

### Identifying TLSs signature genes

TLSs&LAs signature genes were identified from 5 TLSs&LAs+ STs data by differential expression analysis and AUC validation. We first choose four TLSs&LAs + STs data and applying DEGs analysis to identify TLSs&LAs marker genes *FindAllMarkers* function in *Seurat*. Then only common DGEs are further validated in the remain one and genes with average AUC greater than 0.7 in at least 4 samples were keep for further analysis (Supplementary Figure 11A). These analyses resulting in 13 TLSs&LAs sigature genes, including *LTB*, *TRBC2*, *CXCR4*, *CORO1A*, *RPS27A*, *TRAC*, *CCL19*, *TRBC1*, *MS4A1*, *CD79A*, *ZFP36L2*, *CCR7* and *BIRC3*.

### BCR repertoire profiling using bulk RNA-seq

To identify and quantify the IGH and IGL clonotype in bulk RNA-seq samples from Yuan et al. cohort, *MiXCR* (v.3.0.13)^29^ was used to perform this analysis in “shotgun” mode with parameters --species hsa --starting-material rna --only-productive --impute-germline-on-export --assemble “-ObadQualityThreshold=0” --contig-assembly –align “-OsaveOriginalReads=true”. For each clonotype, V gene mutations were extracted from the ‘‘AllVAligments’’ column provided by the *MiXCR* pipeline and summed as described in a previous study^14^.

### Signature gene score analysis and pathway enrichment/activity analysis

The expression activity of gene sets generated in this study (e.g., TLSs&LAs signature genes) or downloaded from other studies (e.g., cytotoxic genes) was quantified by *GSVA* (v.1.34.0)^61^ and *Seurat* function *AddModuleScore* for bulk RNA-seq data and scRNA-seq data, respectively. Immune cell related signature gene sets were collected from previous studies (Supplementary Table 5). DEGs between macrophages from IAC tumor tissues and normal lung tissues were subjected to pathway enrichment analysis using the *Gene Ontology-Biological Processes* (*GOBP*) database and the *enrichr* function from *gseapy* (v.1.0.6)^62^. Pathway activities of epithelial cells were evaluated by *GSVA* with hallmark signature derived from the *MSigDB* (v.7.1)^63^.

## Data availability

The generated STs and scRNA-seq data in this study have been deposited to Genome Sequence Archive (GSA) in BIG Data Center, Beijing Institute of Genomics (BIG) under accession number HRA005812. Processed STs and scRNA-seq data in this study have been deposited in Zenodo (https://doi.org/10.5281/zenodo.8417887). This study additionally included analyses of publicly available datasets. The accession numbers and links of these public datasets are listed in Supplementary Table 10.

## Code availability

The original code used to analyze the data of this study is available at https://github.com/haojiechen94/LUAD_COPs.

## Authors’ contributions

S.Z., Y.S, Y.P., H.C., Y.L and H.H. participated in the study design and study conception. Y.P., P.L., Y.W. and Z.W. obtained patient consent and collected the samples. Y.P. performed single cell RNA sequencing and spatial transcriptomics and processed the data. H.C. and Y.L. performed bioinformatics analyses. Q.Z and Y.P. performed histological evaluations. Y.S. and Y.Z. provided clinical insights. Y.P. and H.C. analyzed and interpreted the data. H.C. and Y.P. conceived the study and wrote the manuscript. S.Z. and Y.S, supervised the study. All authors reviewed and approved the final manuscript.

## Acknowledgement

We sincerely thank Dr Xin Liu from CAS Center for Excellence in Molecular Cell Science for his suggestions on this work.

## Supplementary Figures

**Supplementary Figure 1 Basic quality control Spatial Transcriptomics (STs)**

**(A-B)** Spatial heatmaps of the number of UMIs and features (genes) in each slide and boxplots showing the overall distributions of the number of UMIs and features.

**Supplementary Figure 2 Differentially expressed genes between bulk RNA-seq data from normal lung tissues (bN) and tumor tissues (bT)**

Using DEseq2 to identify significant differentially expressed genes between bN and bT. Those genes with |log2 fold change|>1 and adjusted *p*-value<0.01 were defined as significantly differentially expressed genes.

**Supplementary Figure 3 Evaluating the consistency between transcriptomic signature and histologically annotated areas**

**(A)** HE images and spatial heatmaps of expression of bN and bT signature genes. **(B)** ROC curves for classifying non-cancerous spots (bTAT) in STs data using bN and bT signature genes or randomly selected genes (negative control). P12 was a special case with low AUC and excluded from the downstream analysis. **(C)** Boxplots showing the AUC values in (B).

**Supplementary Figure 4 Quality control metrics across the scRNA-seq dataset**

Statistical summary of cells passing quality control and showing the number of cells, the percentage of UMIs from mitochondrial genes, the number of UMIs, the number of features in each sample.

**Supplementary Figure 5 Dissecting LUAD and normal lung ecosystems by scRNA-seq**

**(A)** UMAP visualization of eight major cell types and cells are colored by their annotated cell types. **(B)** Bubble heatmap showing the percentage of cells expressing major cell type markers (indicated by the size of circle) as well as their relative expression level (indicated by the color of the circle) across all cell types. **(C)** Stacked bar plots showing the fraction of each major cell type derived from each patient (left) and each tissue type (middle). Bar plots showing the absolute number of cells in each major cell type. **(D)** UMAP visualization of night epithelial subclusters and cells are colored by their annotated cell types. **(E)** Bubble heatmap showing the percentage of cells expressing well-known normal epithelial cells markers and malignant cells markers as well as their relative expression level across all epithelial subclusters. **(F)** UMAP visualization of all epithelial cells colored by their inferred CNV scores. Green circle indicates normal epithelial cells and red circle indicates malignant cells. **(G)** The CNV scores of cells from each tissue type.

**Supplementary Figure 6 Workflow of cellular organization patterns identification**

Deconvolution is applied to quantify the cellular composition in each spot and then unsupervised clustering is performed to integrate spots from different slides and identify cellular organization patterns (COPs), COPs are visualized in two dimensional UMAP. Finally, COPs are mapping into the pathologically annotated areas in HE images, by this mean to link the COPs to spatial structures. TAT, tumor adjacent tissues, TLSs, tertiary lymphoid structures, LAs, lymphoid aggregates.

**Supplementary Figure 7 Spatial distribution of COP8 and pathologically annotated papillary tissues in P13**

COP8 colocalized with papillary tissues in P13.

**Supplementary Figure 8 Subclustering of T/NK cells**

**(A)** UMAP visualization of T/NK cells subclusters and cells are colored by their annotated cell types. **(B)** Bubble heatmap showing the percentage of cells expressing T/NK sub cell type/state related markers as well as their relative expression level across all T/NK cells subclusters. (C) Schematic depiction of workflow and specific marker genes used for cell type assignment in T/NK cells subclusters.

**Supplementary Figure 9 Evaluating relative infiltrating levels of different immune cell types in each COP**

**(A)** Cell type specific topic profiles, circle size indicates the specificity of a topic to this cell type. **(B)** The distribution of differences of observed average proportion and permuted average proportion for B cells in each COPs. **(C)** Scaling the mean of each COP distribution in (B) by the standard deviation of the distribution. A larger positive value represents a higher infiltrating level of B cells in this COP. A negative value indicates depletion of B cells in this COP compared with other COPs.

**Supplementary Figure 10 Protumor immune remodeling is also evidenced in scRNA-seq data**

**(A-D)** Box plots showing the percentages of functional or dysfunctional immune cell type (group) across different tissue types. **(E)** Line plots showing the changes in percentages among cell type (group) in (A-D) across different tissue types.

**Supplementary Figure 11 Cytotoxic activity of CD4+ T cell subclusters**

**(A)** Bubble heatmap showing the percentage of cells expressing cytotoxic genes as well as their relative expression levels in each CD4+ T cell subpopulation. (B) UMAP plot visualization of CD4+ T cells colored by annotated cell types (left) and cytotoxic genes scores (right).

**Supplementary Figure 12 Schematic of TLSs&LAs signature genes identification and its association with ICI response**

**(A)** Workflow of TLSs&LAs signature genes identification, choosing four TLSs&LAs + STs data and applying DEGs analysis to identify TLSs&LAs marker genes, only common DGEs are further validated in the remain one. **(B)** Boxplots of TLSs&LAs signature scores in DB and NDB samples in *Hwang et al.* cohort. DB, durable benefit, NDB, non-durable benefit.

**Supplementary Figure 13 Location and proportion of B cells and plasma cells in STs data from TLSs&LAs+ samples or TLSs&LAs-samples**

HE images showing the locations of TLSs and LAs spots, spatial heatmap showing the location and proportion of B cells and plasma cells.

**Supplementary Figure 14 Molecular and function potential alterations of macrophages from different tissue types**

**(A)** Volcano plot showing the DEGs between monocyte-derived macrophages (MDMs) from IAC tumor tissues and normal lung tissues. Antigen presenting related molecules (MHC genes) are highlighted in green. M2 polarization related molecules are highlighted in red. **(B-C)** Violin plots of signature scores of MHC II molecules (B), oxidative phosphorylation pathway (C) and M2 polarization associated genes (D), across MDMs from different tissue types.

**Supplementary Figure 15 Increased number of cell-to-cell interactions during pathological progression**

The number of *CellphoneDB* inferred cell-to-cell interactions between normal epithelial cells/MIA malignant cells/IAC malignant cells and other cell types in ecosystem.

**Supplementary Figure 16 Identification of malignant cells and T cells in a public LUAD scRNA-seq dataset including tumor tissues from AIS, MIA and IAC**

A total of 38 cell subpopulations were identified through clustering, and each subpopulation was characterized based on the expression of specific marker genes. AT1 cells were identified as subpopulations that exhibited high expression of both *EPCAM* and *AGER*. Ciliated epithelial cells were defined as subpopulations with high expression of both *EPCAM* and *FOXJ1*. AT2 cells were determined as subpopulations showing high expression of both *EPCAM* and *SFTPC*. Subpopulations that expressed only *EPCAM* without the usual marker genes associated with AT1, ciliated epithelial cells, or AT2 cells were classified as malignant cells. Additionally, T cells were identified as subpopulations with high expression of both *CD3D* and *CD3E*.

**Supplementary Figure 17 Higher CNV scores in PC4 and PC4 malignant cells from IAC**

**(A-B)** Box plots illustrating the CNV scores for each phenotypic clusters (A) and PC4 in MIA and IAC (B).

## Supplementary Tables

**Supplementary Table 1 Clinical information of patients in this study**

**Supplementary Table 2 Marker genes used to define major cell types and related references**

**Supplementary Table 3 Marker genes used to define normal epithelial cells and related references**

**Supplementary Table 4 Marker genes used to define immune cells and related references Supplementary Table 5 Immune cell related signature gene sets in this study**

**Supplementary Table 6 TLSs/LAs signature defined by performing leave-one-out cross validation on TLSs/LAs+ STs samples**

**Supplementary Table 7 Differentially expressed genes between IAC and normal lung tissues infiltrated Monocyte-derived macrophages**

**Supplementary Table 8 Signature genes of phenotypic cluster 4 (PC4) Supplementary Table 9 Predicted sensitive drugs of PC4**

**Supplementary Table 10 Accession numbers for public datasets used in this study**

