## Supplementary Figures for "Spatiotemporal Profiling Unveiling the Cellular Organization Patterns and Local Protumoral Immune Microenvironment Remodeling in Early Lung Adenocarcinoma Progression"

### Supplementary Figure1

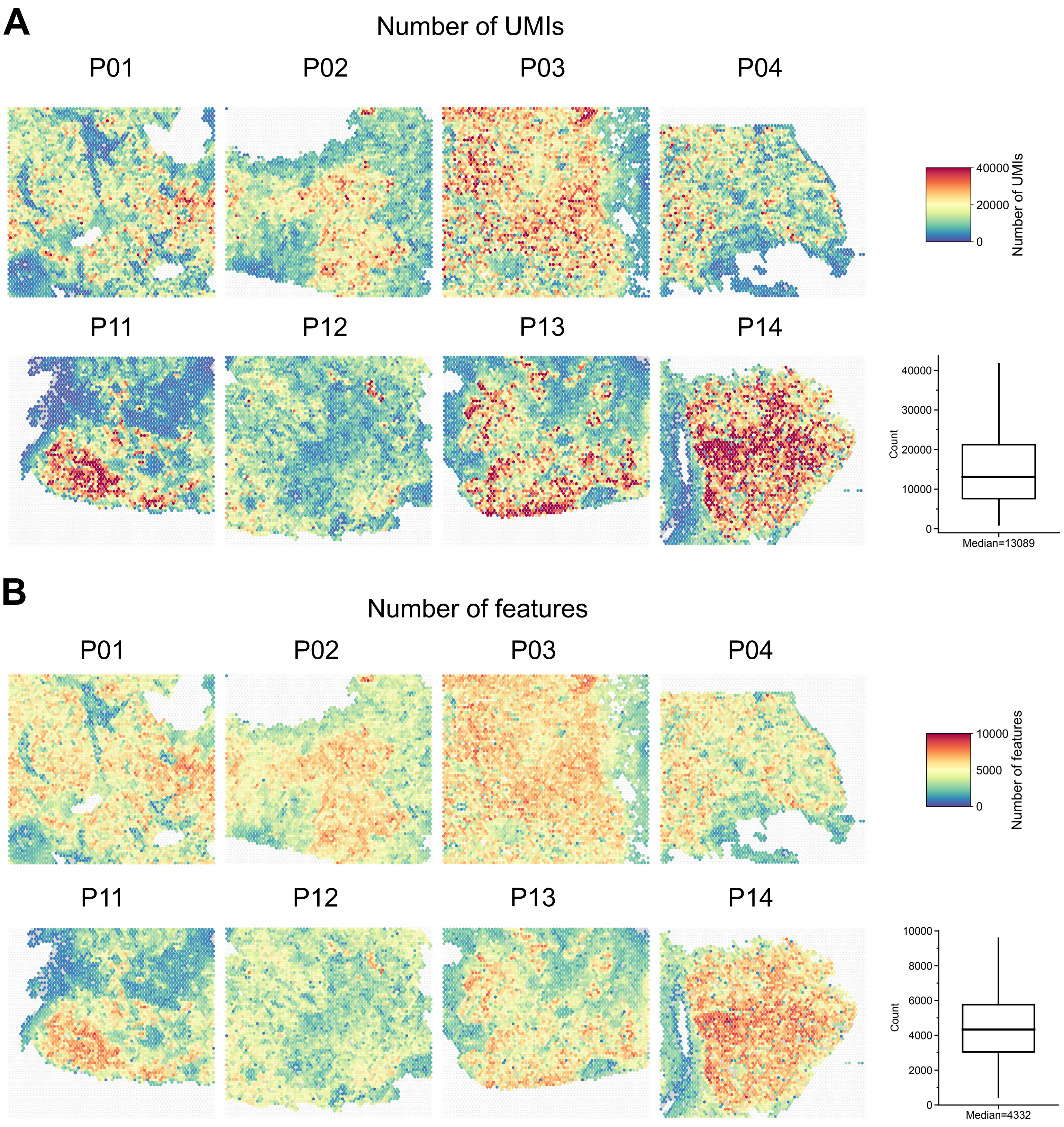

**Supplementary Figure 1 Basic quality control Spatial Transcriptomics (STs)**

**(A-B)** Spatial heatmaps of the number of UMIs and features (genes) in each slide and boxplots showing the overall distributions of the number of UMIs and features.

### Supplementary Figure2

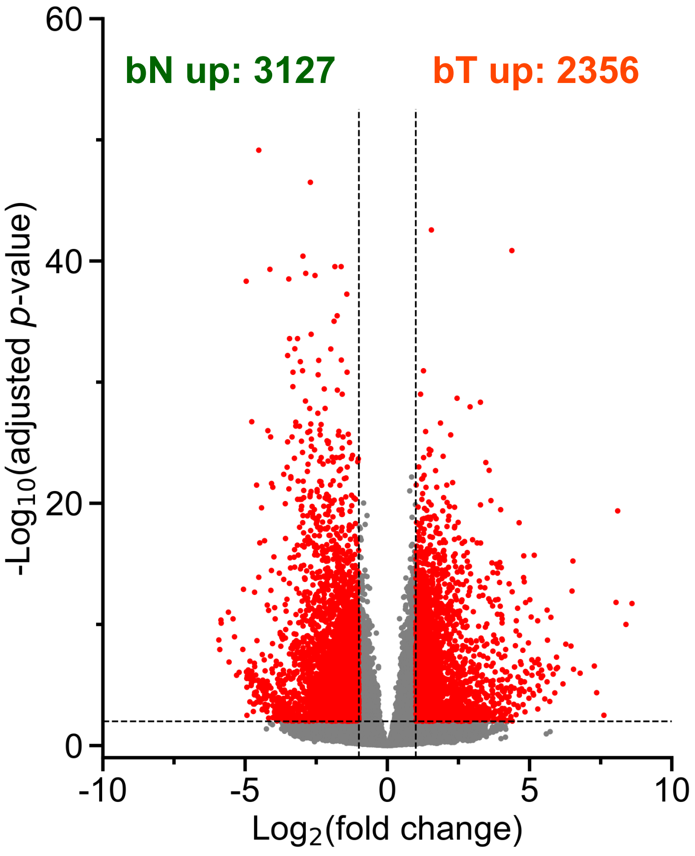

**Supplementary Figure 2 Differentially expressed genes between bulk RNA-seq data from normal lung tissues (bN) and tumor tissues (bT)**

Using DEseq2 to identify significant differentially expressed genes between bN and bT. Those genes with |log2 fold change|>1 and adjusted *p*-value<0.01 were defined as significantly differentially expressed genes.

### Supplementary Figure 3

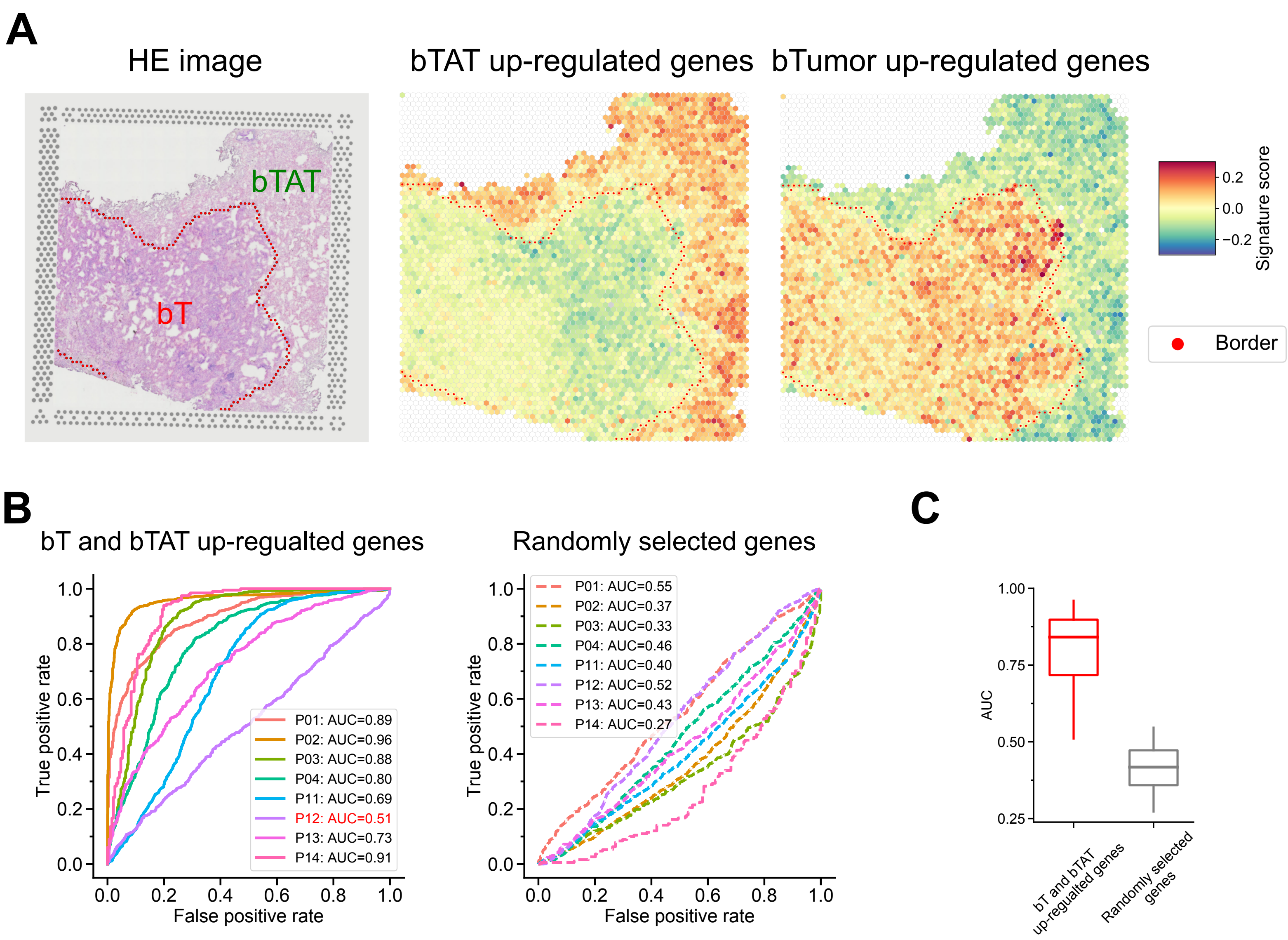

**Supplementary Figure 3 Evaluating the consistency between transcriptomic signature and histologically annotated areas**

**(A)** HE images and spatial heatmaps of expression of bN and bT signature genes. **(B)** ROC curves for classifying non-cancerous spots (bTAT) in STs data using bN and bT signature genes or randomly selected genes (negative control). P12 was a special case with low AUC and excluded from the downstream analysis. **(C)** Boxplots showing the AUC values in (B).

### Supplementary Figure 4

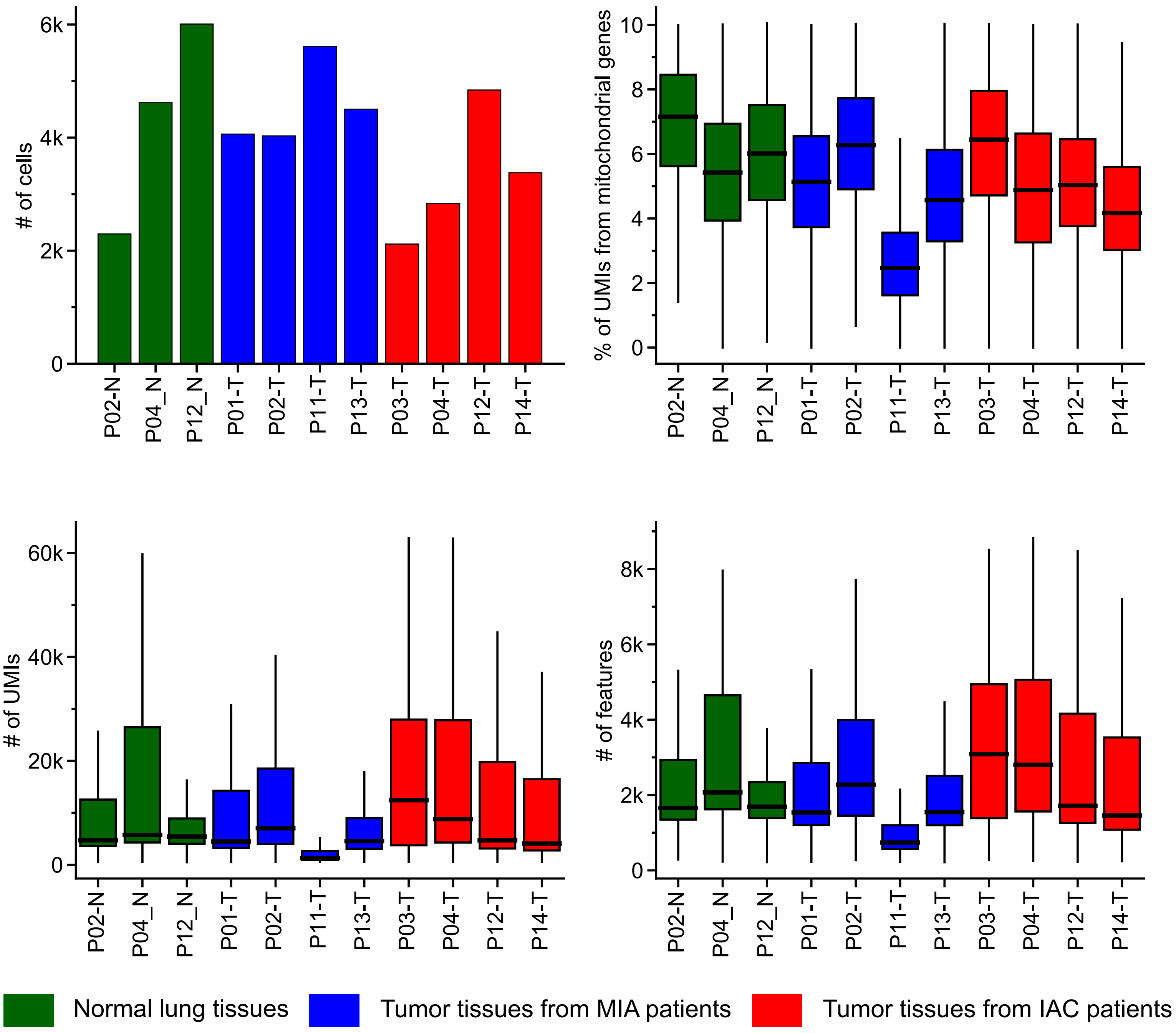

**Supplementary Figure 4 Quality control metrics across the scRNA-seq dataset**

Statistical summary of cells passing quality control and showing the number of cells, the percentage of UMIs from mitochondrial genes, the number of UMIs, the number of features in each sample.

### Supplementary Figure 5

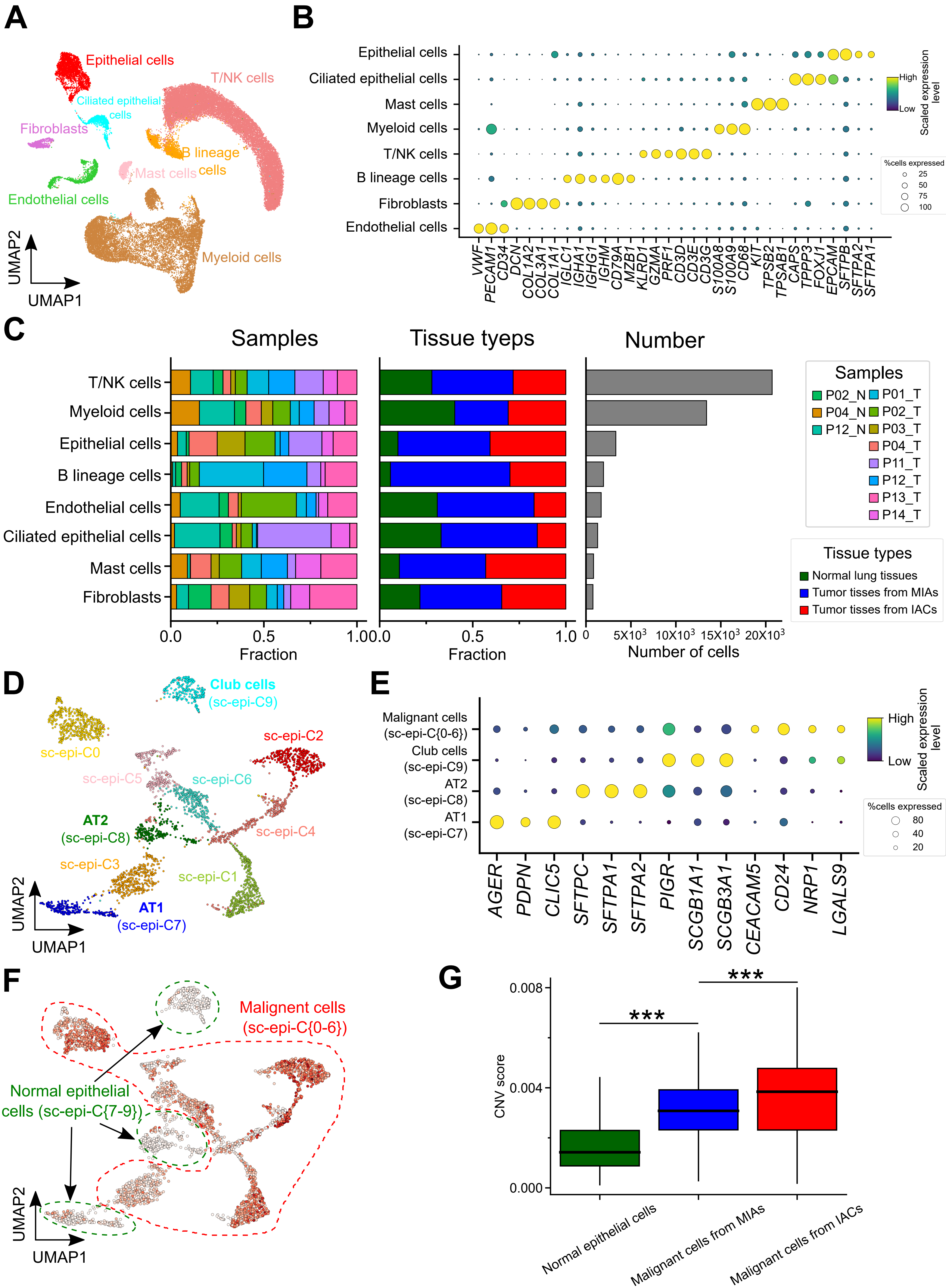

**Supplementary Figure 5 Dissecting LUAD and normal lung ecosystems by scRNA-seq**

**(A)** UMAP visualization of eight major cell types and cells are colored by their annotated cell types. **(B)** Bubble heatmap showing the percentage of cells expressing major cell type markers (indicated by the size of circle) as well as their relative expression level (indicated by the color of the circle) across all cell types. **(C)** Stacked bar plots showing the fraction of each major cell type derived from each patient (left) and each tissue type (middle). Bar plots showing the absolute number of cells in each major cell type. **(D)** UMAP visualization of night epithelial subclusters and cells are colored by their annotated cell types. **(E)** Bubble heatmap showing the percentage of cells expressing well-known normal epithelial cells markers and malignant cells markers as well as their relative expression level across all epithelial subclusters. **(F)** UMAP visualization of all epithelial cells colored by their inferred CNV scores. Green circle indicates normal epithelial cells and red circle indicates malignant cells. **(G)** The CNV scores of cells from each tissue type.

### Supplementary Figure 6

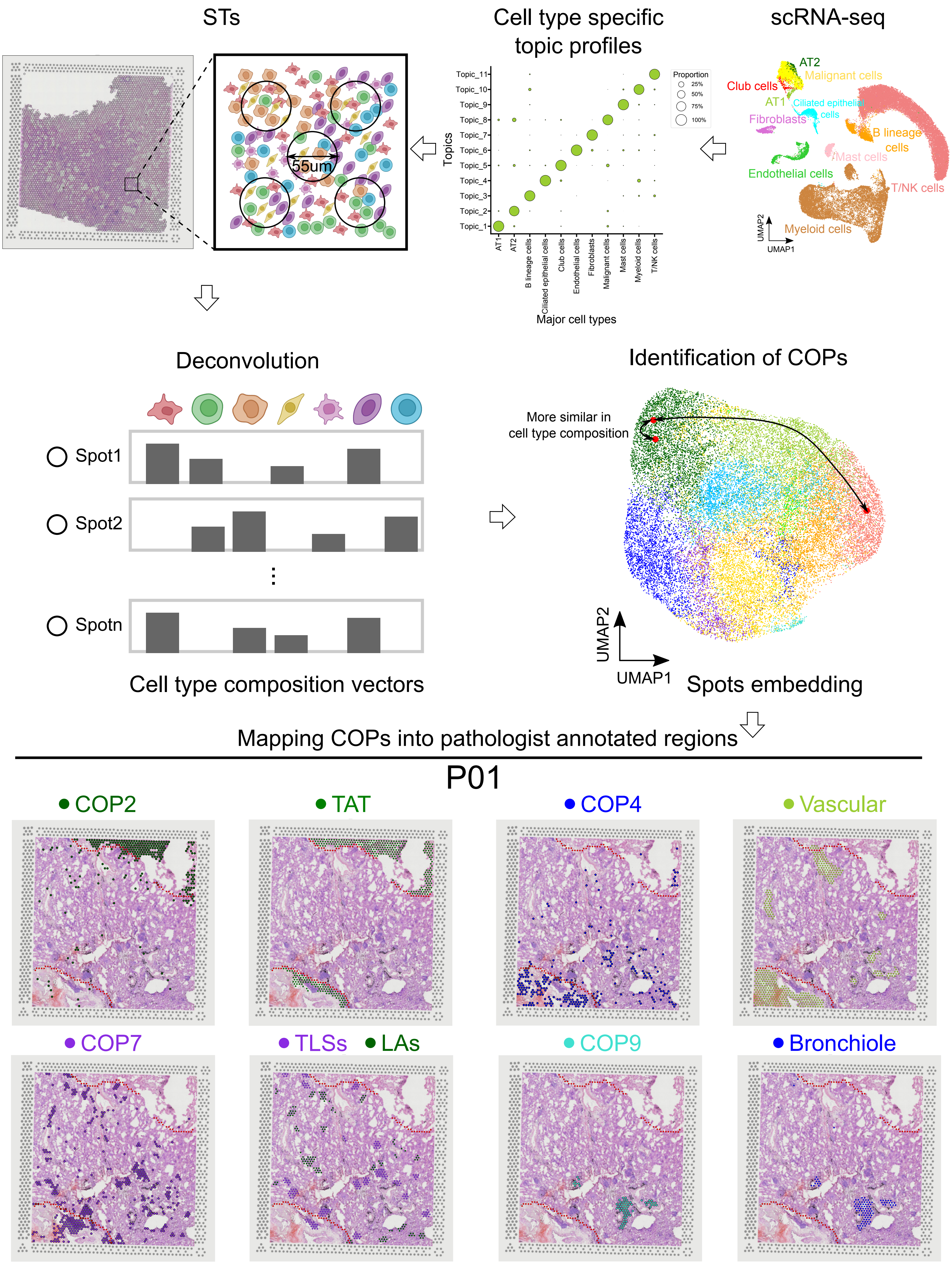

**Supplementary Figure 6 Workflow of cellular organization patterns identification**

Deconvolution is applied to quantify the cellular composition in each spot and then unsupervised clustering is performed to integrate spots from different slides and identify cellular organization patterns (COPs), COPs are visualized in two dimensional UMAP. Finally, COPs are mapping into the pathologically annotated areas in HE images, by this mean to link the COPs to spatial structures. TAT, tumor adjacent tissues, TLSs, tertiary lymphoid structures, LAs, lymphoid aggregates.

### Supplementary Figure 7

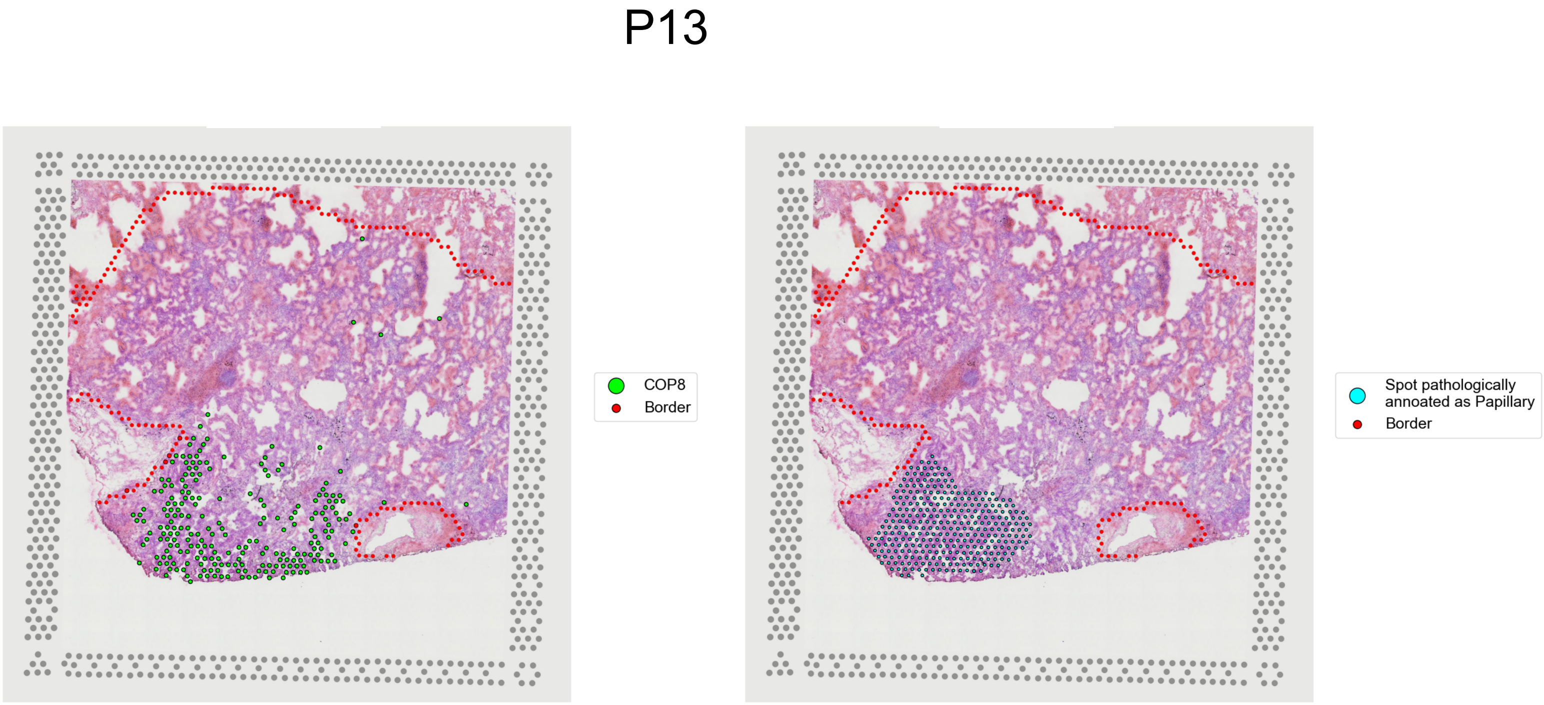

**Supplementary Figure 7 Spatial distribution of COP8 and pathologically annotated papillary tissues in P13**

COP8 colocalized with papillary tissues in P13.

### Supplementary Figure 8

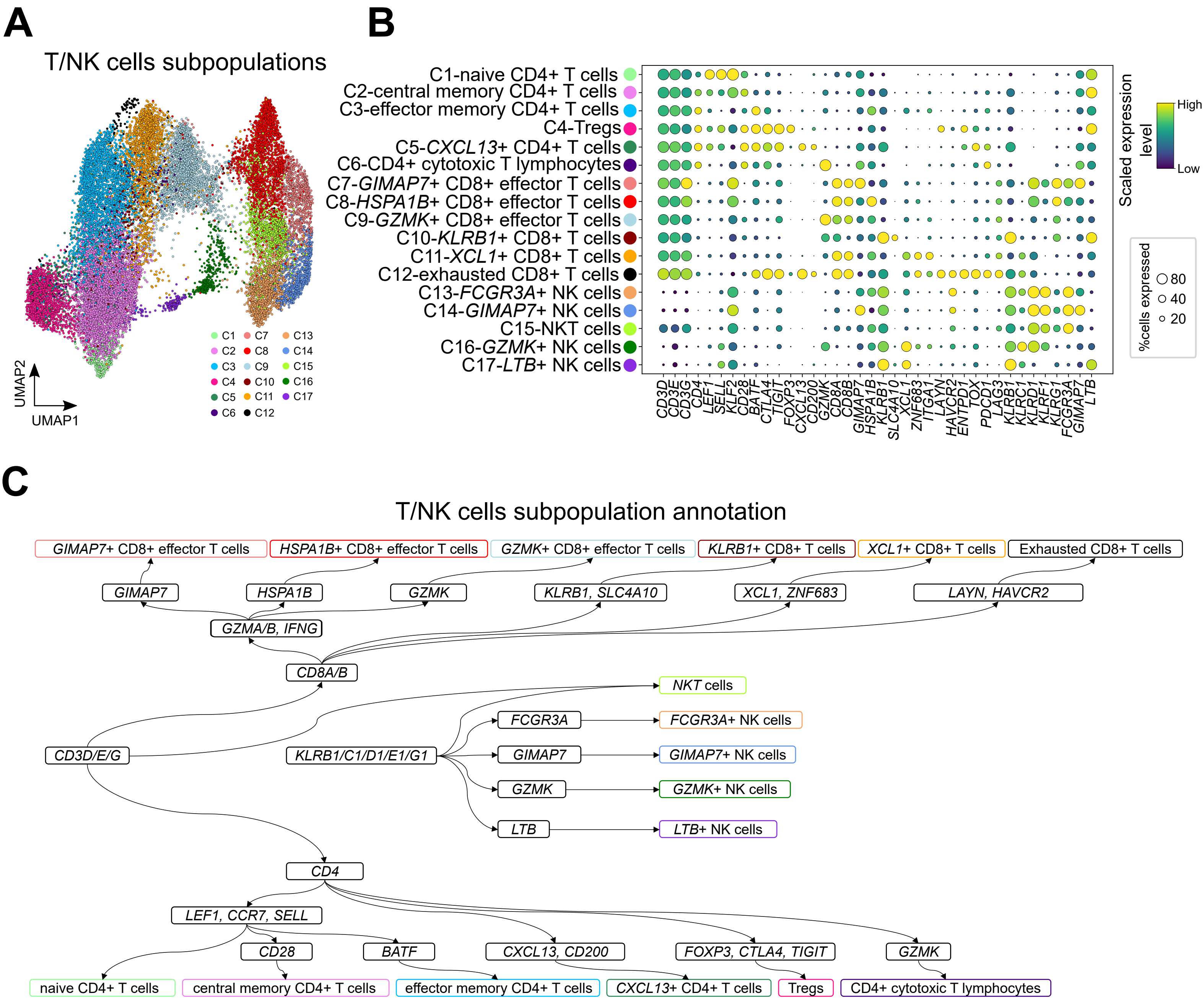

**Supplementary Figure 8 Subclustering of T/NK cells**

**(A)** UMAP visualization of T/NK cells subclusters and cells are colored by their annotated cell types. **(B)** Bubble heatmap showing the percentage of cells expressing T/NK sub cell type/state related markers as well as their relative expression level across all T/NK cells subclusters. **(C)** Schematic depiction of workflow and specific marker genes used for cell type assignment in T/NK cells subclusters.

### Supplementaty Figure 9

**A**

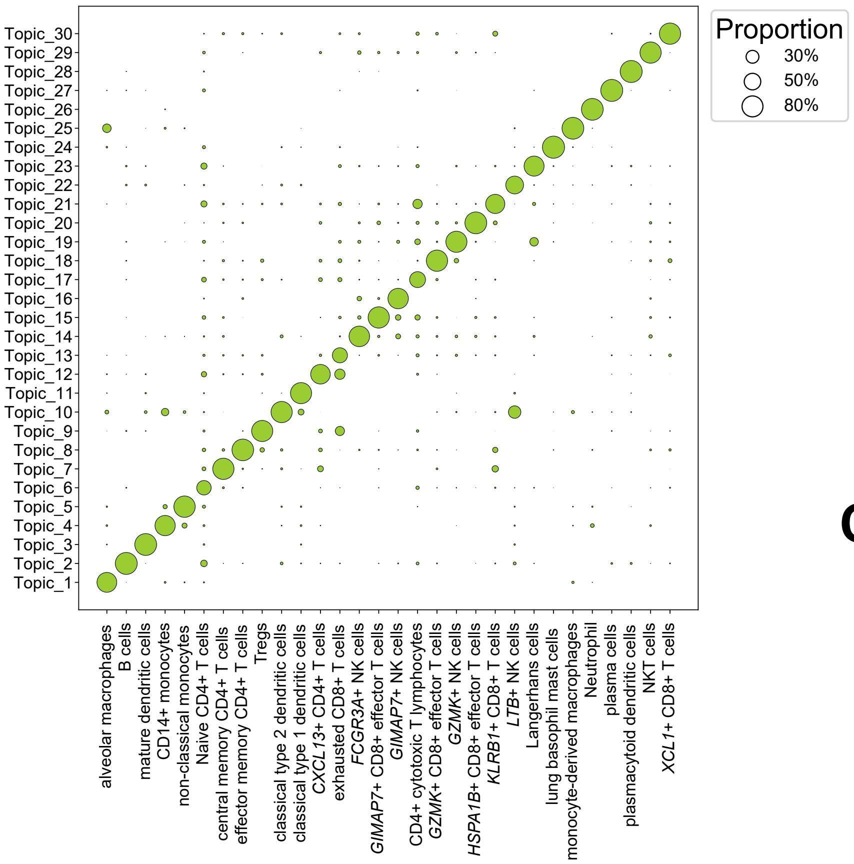


**B**

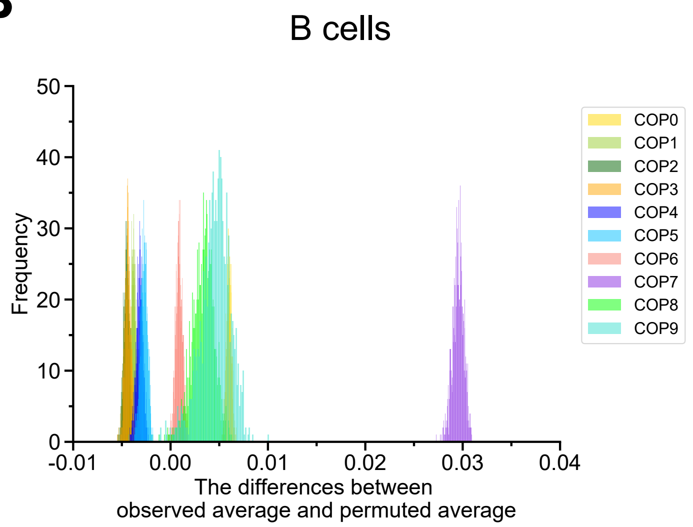


**C**

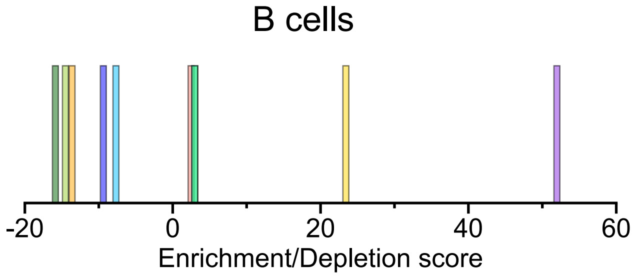

**Supplementary Figure 9 Evaluating relative infiltrating levels of different immune cell types in each COP**

**(A)** Cell type specific topic profiles, circle size indicates the specificity of a topic to this cell type. **(B)** The distribution of differences of observed average proportion and permuted average proportion for B cells in each COPs. **(C)** Scaling the mean of each COP distribution in (B) by the standard deviation of the distribution. A larger positive value represents a higher infiltrating level of B cells in this COP. A negative value indicates depletion of B cells in this COP compared with other COPs.

### Supplementary Figure 10

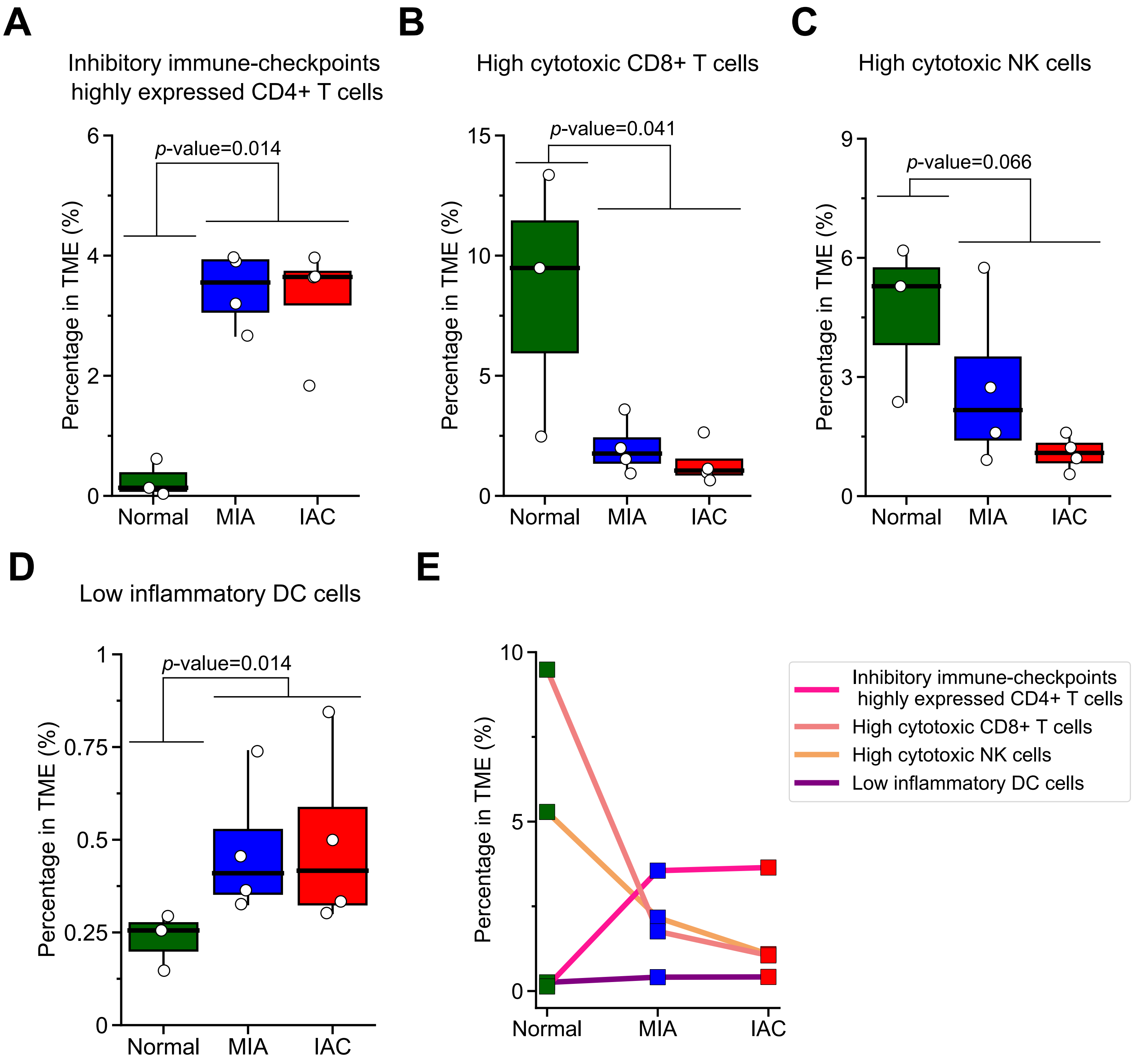

**Supplementary Figure 10 Protumor immune remodeling is also evidenced in scRNA-seq data**

**(A-D)** Box plots showing the percentages of functional or dysfunctional immune cell type (group) across different tissue types. **(E)** Line plots showing the changes in percentages among cell type (group) in (A-D) across different tissue types.

### Supplementary Figure 11

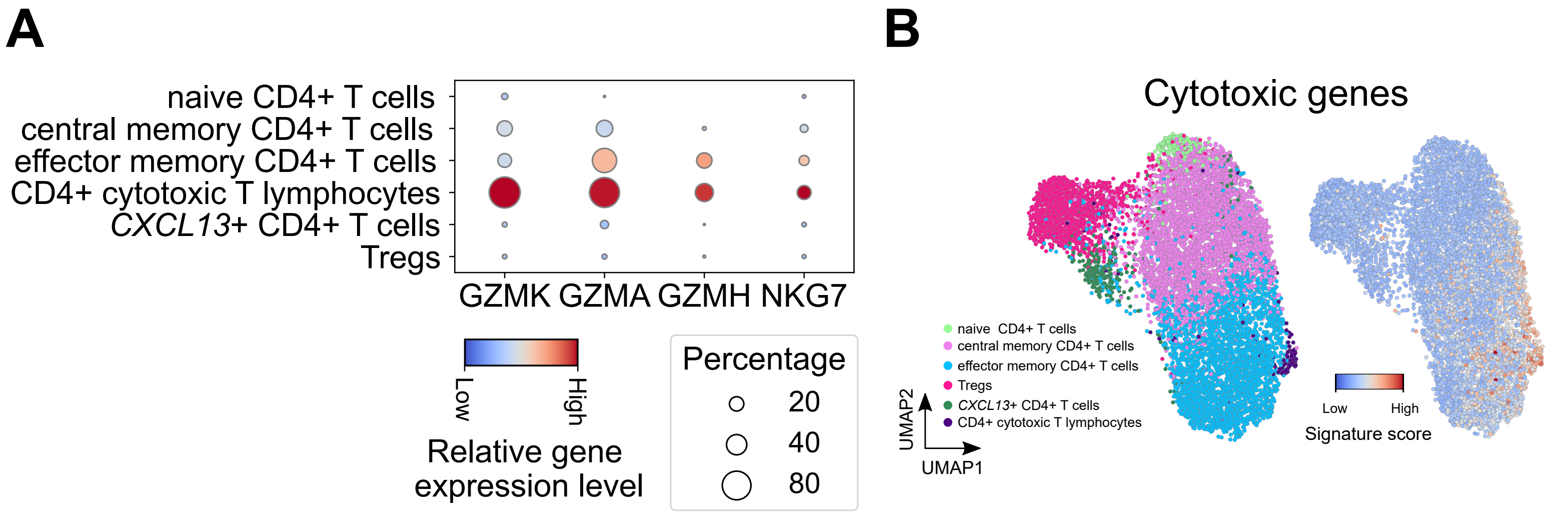

**Supplementary Figure 11 Cytotoxic activity of CD4+ T cell subclusters**

**(A)** Bubble heatmap showing the percentage of cells expressing cytotoxic genes as well as their relative expression levels in each CD4+ T cell subpopulation. (B) UMAP plot visualization of CD4+ T cells colored by annotated cell types (left) and cytotoxic genes scores (right).

### Supplementary Figure 12

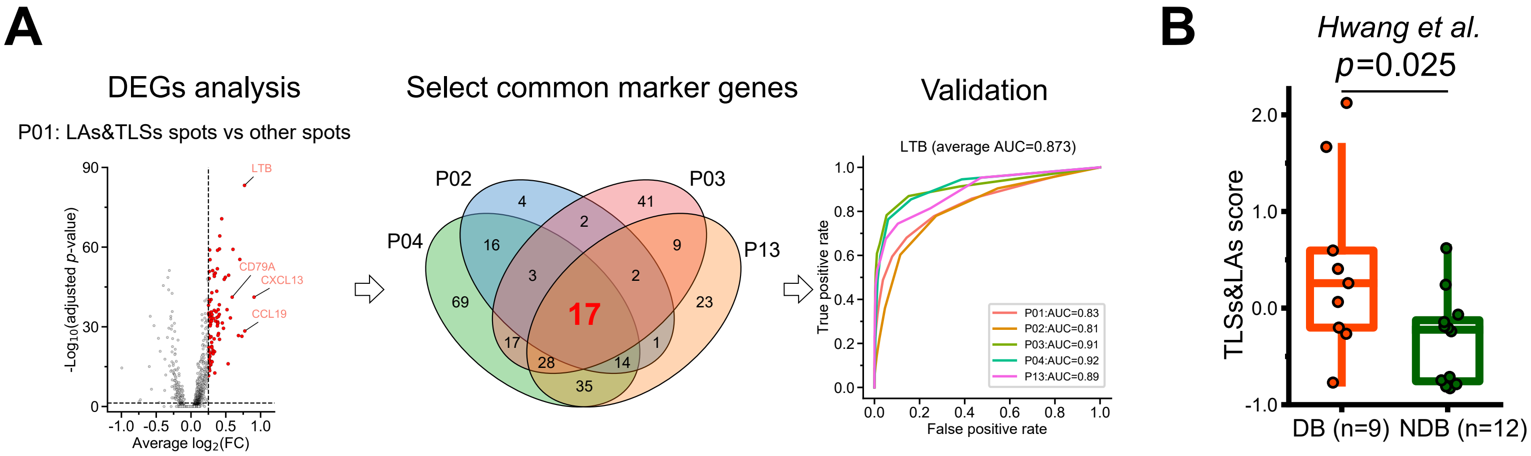

**Supplementary Figure 12 Schematic of TLSs&LAs signature genes identification and its association with ICI response**

**(A)** Workflow of TLSs&LAs signature genes identification, choosing four TLSs&LAs + STs data and applying DEGs analysis to identify TLSs&LAs marker genes, only common DGEs are further validated in the remain one. **(B)** Boxplots of TLSs&LAs signature scores in DB and NDB samples in *Hwang et al.* cohort. DB, durable benefit, NDB, non-durable benefit.

### Supplementary Figure 13

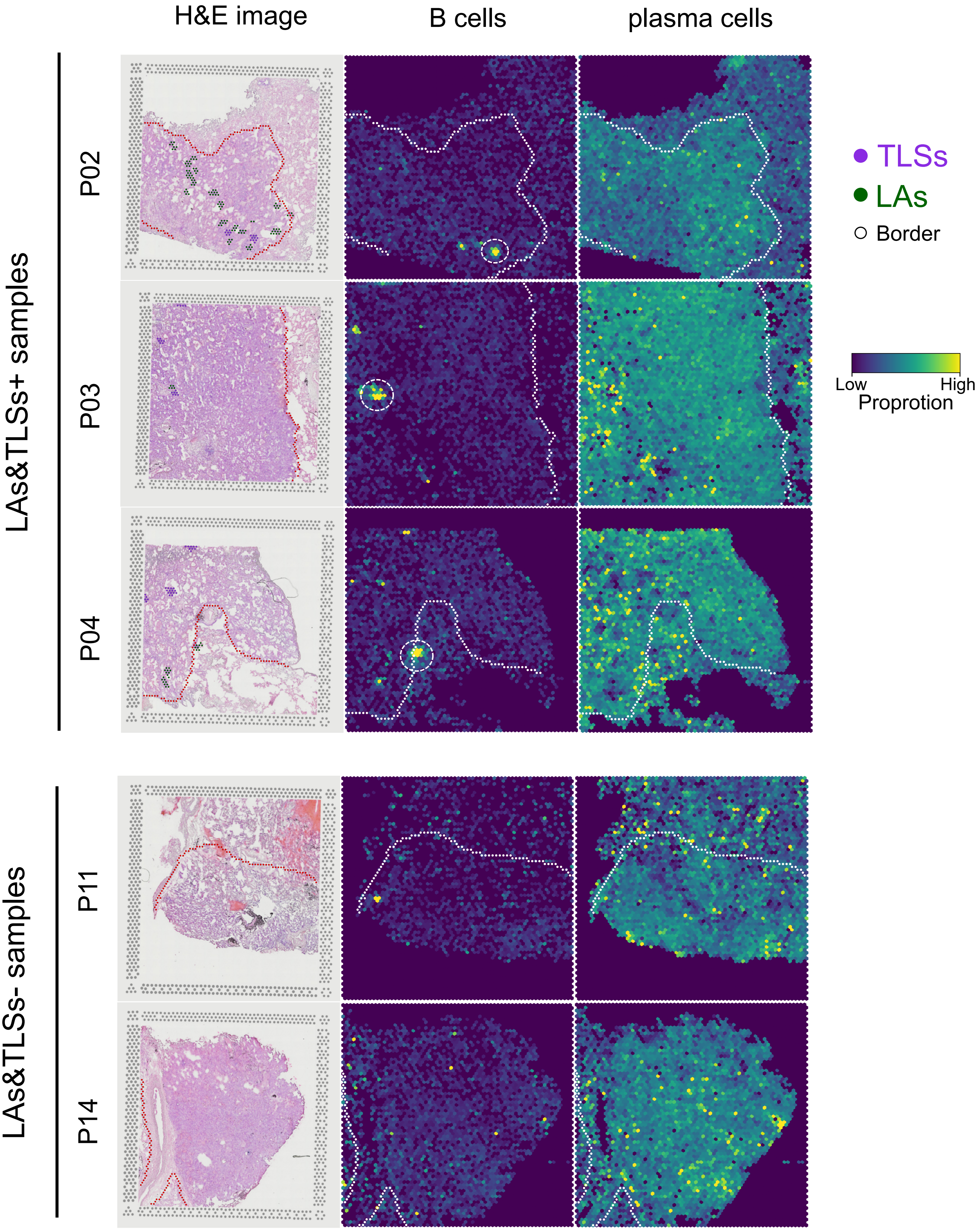

**Supplementary Figure 13 Location and proportion of B cells and plasma cells in STs data from TLSs&LAs+ samples or TLSs&LAs- samples**

HE images showing the locations of TLSs and LAs spots, spatial heatmap showing the location and proportion of B cells and plasma cells.

### Supplementary Figure 14

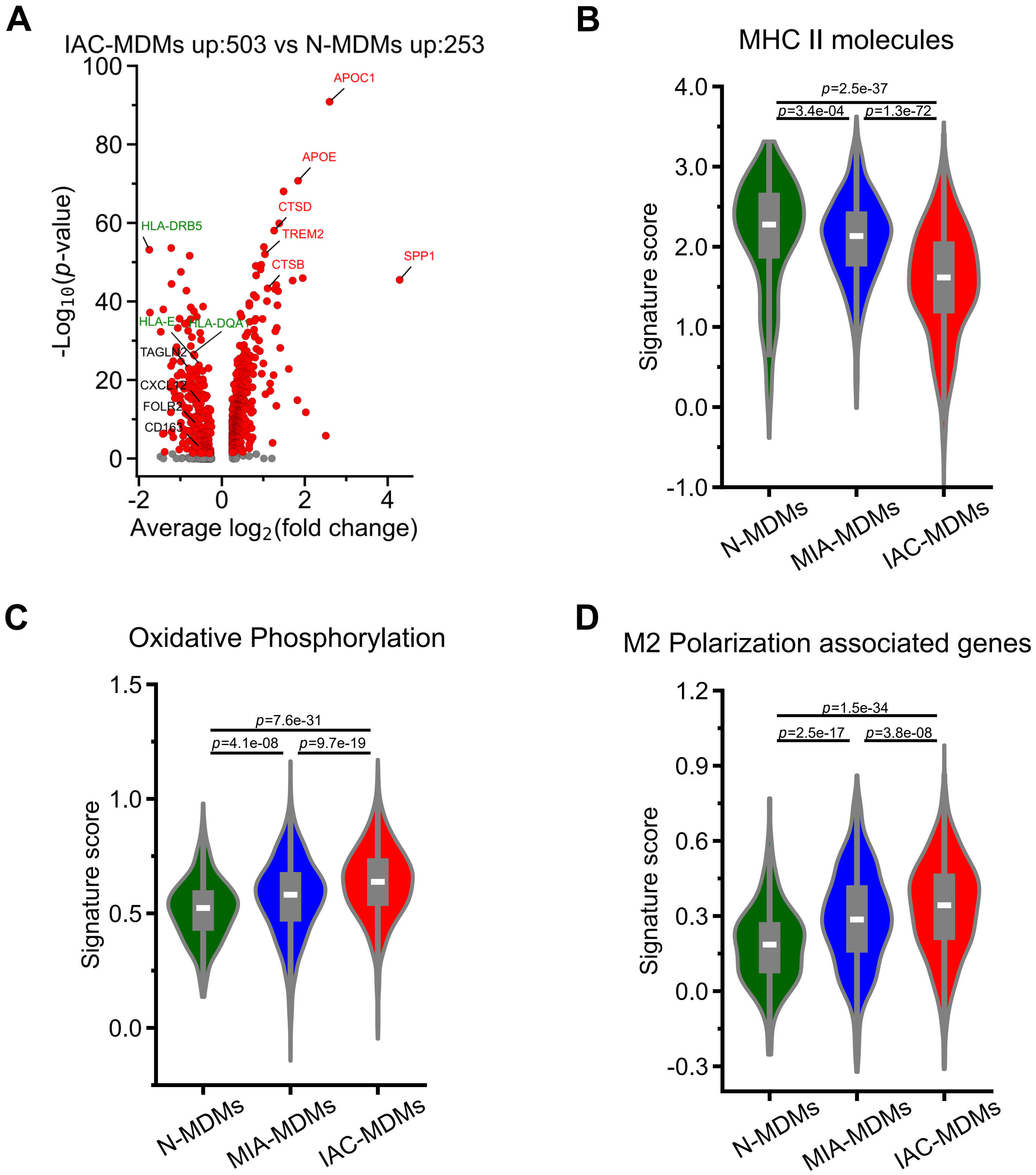

**Supplementary Figure 14 Molecular and function potential alterations of macrophages from different tissue types**

**(A)** Volcano plot showing the DEGs between monocyte-derived macrophages (MDMs) from IAC tumor tissues and normal lung tissues. Antigen presenting related molecules (MHC genes) are highlighted in green. M2 polarization related molecules are highlighted in red. **(B-C)** Violin plots of signature scores of MHC II molecules (B), oxidative phosphorylation pathway (C) and M2 polarization associated genes (D), across MDMs from different tissue types.

### Supplementary Figure 15

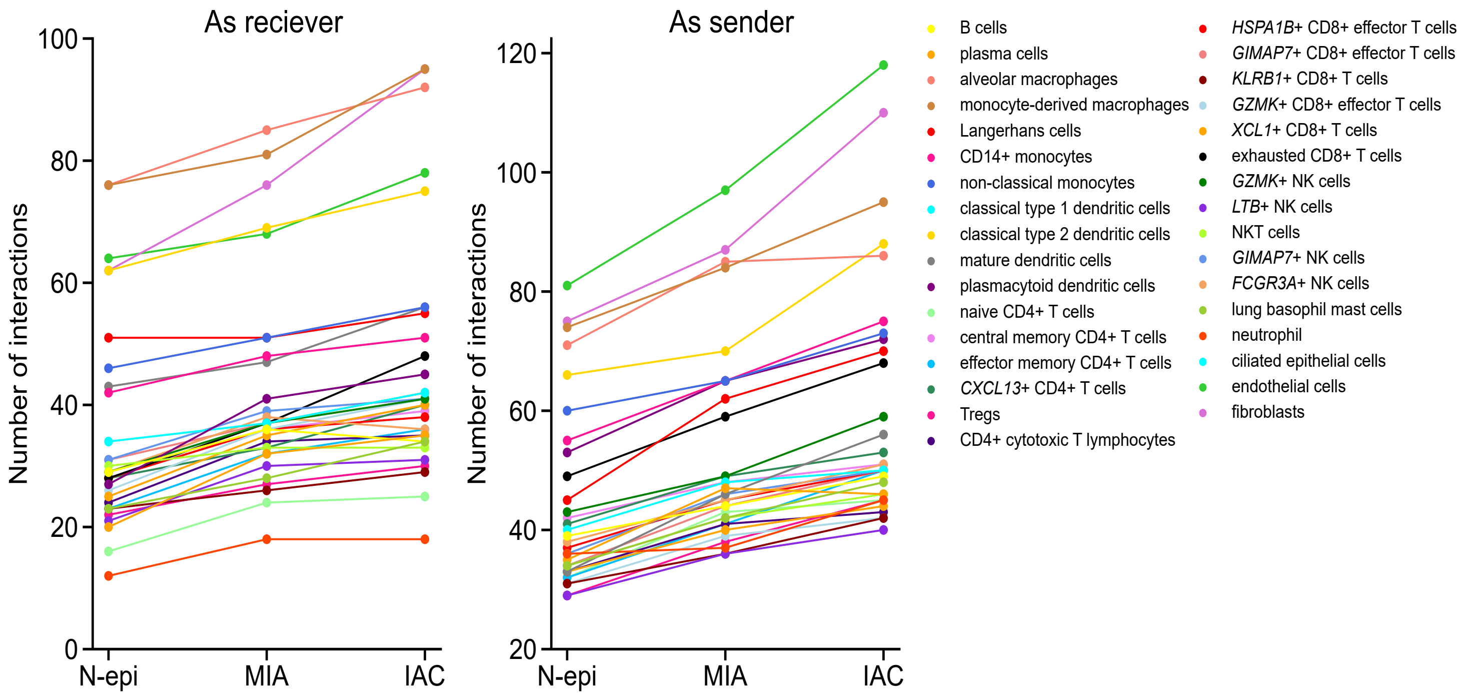

**Supplementary Figure 15 Increased number of cell-to-cell interactions during pathological progression**

The number of *CellphoneDB* inferred cell-to-cell interactions between normal epithelial cells/MIA malignant cells/IAC malignant cells and other cell types in ecosystem.

### Supplementary Figure 16

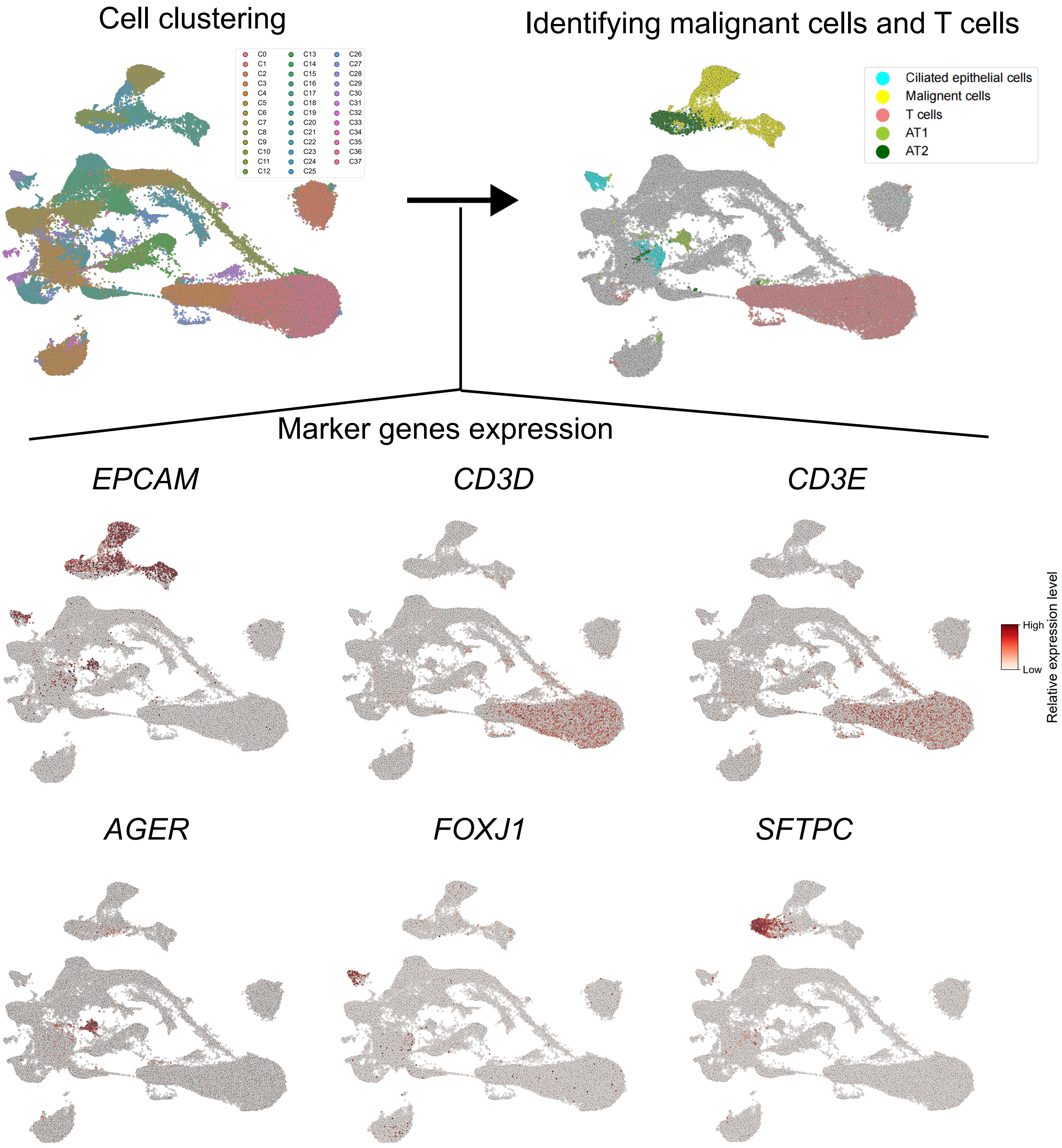

**Supplementary Figure 16 Identification of malignant cells and T cells in a public LUAD scRNA-seq dataset including tumor tissues from AIS, MIA and IAC**

A total of 38 cell subpopulations were identified through clustering, and each subpopulation was characterized based on the expression of specific marker genes. AT1 cells were identified as subpopulations that exhibited high expression of both *EPCAM* and *AGER*. Ciliated epithelial cells were defined as subpopulations with high expression of both *EPCAM* and *FOXJ1*. AT2 cells were determined as subpopulations showing high expression of both *EPCAM* and *SFTPC*. Subpopulations that expressed only *EPCAM* without the usual marker genes associated with AT1, ciliated epithelial cells, or AT2 cells were classified as malignant cells. Additionally, T cells were identified as subpopulations with high expression of both *CD3D* and *CD3E*.

#### Supplementary Figure 17

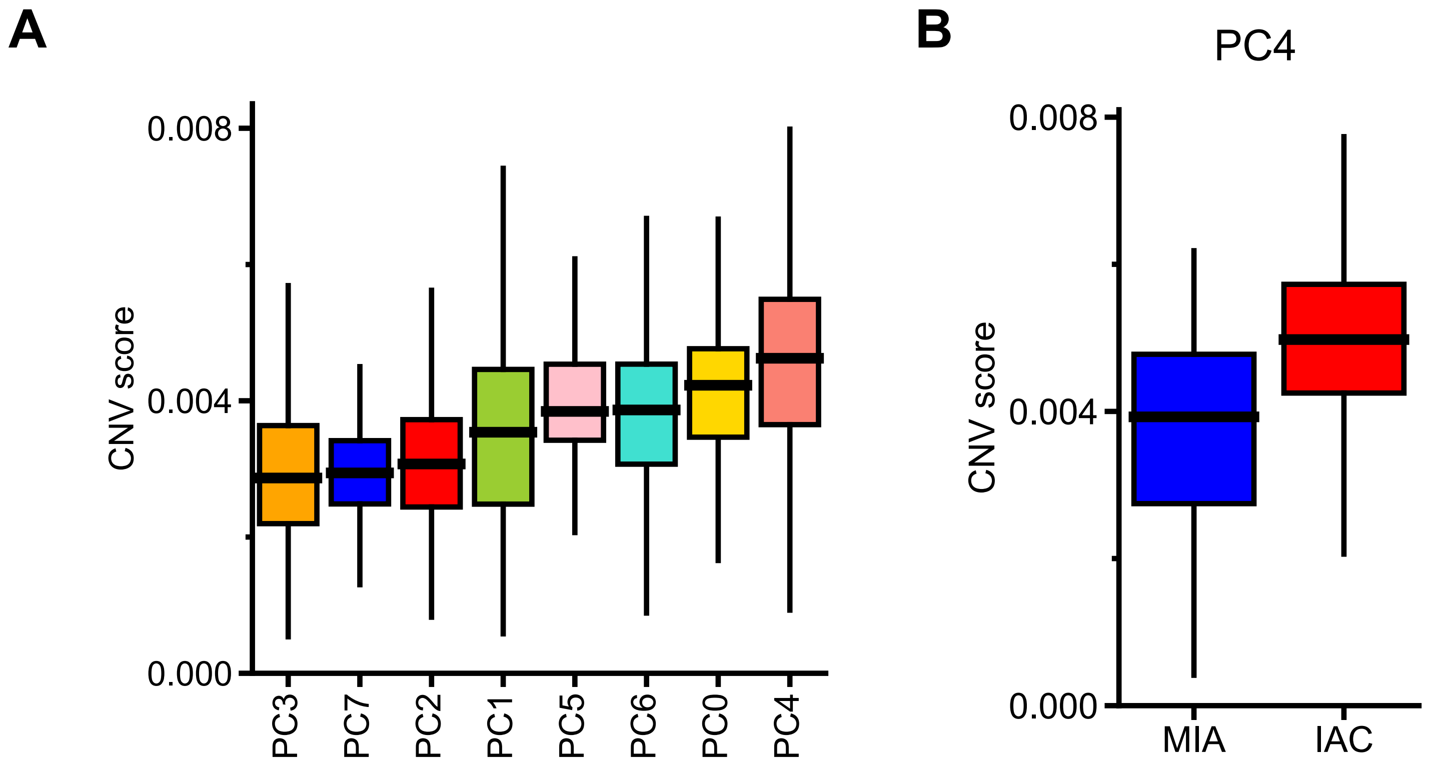

**Supplementary Figure 17 Higher CNV scores in PC4 and PC4 malignant cells from IAC**

**(A-B)** Box plots illustrating the CNV scores for each phenotypic clusters (A) and PC4 in MIA and IAC (B).
